# Antagonistic modules, SIB1 and LSD1, regulate photosynthesis-associated nuclear genes via GOLDEN2-LIKE transcription factors in Arabidopsis

**DOI:** 10.1101/2020.11.22.393603

**Authors:** Mengping Li, Keun Pyo Lee, Tong Liu, Dogra Vivek, Jianli Duan, Mengshuang Li, Weiman Xing, Chanhong Kim

**Author notes:** These authors contributed equally to this work (Mengping Li, Keun Pyo Lee). Biotechnology Division, CSIR-Institute of Himalayan Bioresource Technology, Palampur 176061, India.

## Abstract

GOLDEN2-LIKE (GLK) transcription factors drive the expression of photosynthesis-associated nuclear genes (PhANGs), indispensable for chloroplast biogenesis. We previously demonstrated that the salicylic acid (SA)-induced SIGMA FACTOR-BINDING PROTEIN 1 (SIB1), a transcription coregulator and positive regulator of cell death, interacts with GLK1 and GLK2 to reinforce their activities. The SIB1-GLK interaction raises the level of light-harvesting antenna proteins in photosystem II, aggravating photoinhibition and singlet oxygen (^1^O_2_) burst. ^1^O_2_ then contributes to SA-induced cell death via EXECUTER 1 (EX1, ^1^O_2_ sensor protein)-mediated retrograde signaling upon reaching a critical level. We now reveal that LESION-SIMULATING DISEASE 1 (LSD1), a transcription coregulator and negative regulator of SA-primed cell death, interacts with GLK1/2 to repress their activities. Consistently, the overexpression of LSD1 represses GLK target genes including PhANGs, whereas the loss of LSD1 increases their expression. Remarkably, LSD1 overexpression inhibits chloroplast biogenesis, resembling the characteristic *glk1glk2* double mutant phenotype. The subsequent chromatin immunoprecipitation analysis coupled with quantitative PCR further reveals that LSD1 inhibits the DNA-binding activity of GLK1 towards its target promoters. The SA-induced nuclear-targeted SIB1 appears to counteractively interact with GLK1/2, leading to the activation of EX1-mediated ^1^O_2_ signaling. Taken together, we provide a working model that SIB1 and LSD1, mutually exclusive SA-signaling components, antagonistically regulate GLK1/2 to fine-tune the expression of PhANGs, thereby modulating ^1^O_2_ homeostasis and related stress responses.

## INTRODUCTION

Chloroplasts communicate with the nucleus via retrograde signaling (RS) in response to the ever-changing environment. Upon exposure to unfavorable environmental conditions, chloroplasts downregulate photosynthesis-associated nuclear genes (PhANGs), referred to as biogenic RS, but stimulate the expression of stress-related genes via alternate RS pathways, collectively called operational RS. The nuclear-encoded chloroplast GENOMES UNCOUPLED 1 (GUN1) protein plays a pivotal role in the biogenic RS (Nott et al., 2006). GUN1 integrates various retrograde signals released by the disturbance in plastid gene expression, redox homeostasis, and tetrapyrrole biosynthesis in chloroplasts (Chan et al., 2016; Koussevitzky et al., 2007; Nott et al., 2006). The well-known downstream targets of GUN1-mediated RS are two nuclear genes encoding the GOLDEN2-LIKE (GLK) transcription factors (TFs) (Martin et al., 2016; Waters et al., 2009). In fact, GUN1-mediated RS represses *GLK* transcription. In *Arabidopsis thaliana* (Arabidopsis), GLK1 and GLK2 function redundantly to express PhANGs, promoting chloroplast biogenesis. Consistently, the loss of both GLKs significantly impairs chloroplast biogenesis (Fitter et al., 2002).

Recent studies discovered an unexpected function of GLKs towards plant immune responses. The steady-state levels of salicylic acid (SA)-responsive genes are significantly lower in GLK1-overexpressing (*oxGLK1*) Arabidopsis transgenic plants relative to wild-type (WT) plants (Savitch et al., 2007). Accordingly, the *oxGLK1* plants are susceptible to the biotrophic pathogen *Hyaloperonospora arabidopsidis* (*Hpa*) Noco2, while *glk1 glk2* double knockout mutant plants are more resistant compared to WT plants (Murmu et al., 2014). However, other studies reported that GLKs confer resistance towards the cereal fungal pathogen *Fusarium graminearum* (Savitch et al., 2007), necrotrophic fungal pathogen *Botrytis cinerea* (Murmu et al., 2014), and the Cucumber mosaic virus (Han et al., 2016). These findings indicate that multiple regulatory circuits (positive and negative) may differently modulate GLK activity towards various microbial pathogens.

We lately demonstrated that the nuclear-targeted SIGMA FACTOR-BINDING PROTEIN 1 (SIB1), a defense-related transcription coregulator, interacts with GLK1/2 in response to an increase in foliar SA (Lai et al., 2011; Lv et al., 2019). In Arabidopsis *lesion-simulating disease 1* (*lsd1*) mutant grown under continuous light (CL) conditions, the transiently increased level of SA rapidly induces the otherwise undetectable SIB1, leading to its accumulation in both the nucleus and the chloroplasts (Lai et al., 2011; Lv et al., 2019). It is important to note that the extended daylength is one of the lesion-triggering external factors evoking SA-dependent runaway (uncontrolled) cell death (RCD) in the *lsd1* mutant (Dietrich et al., 1994; Lv et al., 2019). The SA receptor Nonexpresser of PR genes 1 (NPR1) induces the expression of *SIB1* and the dual targeting of SIB1 also occurs in WT plants after SA treatment (Lai et al., 2011; Lv et al., 2019; Xie et al., 2010). Whereas the loss of NPR1 abolishes *lsd1* RCD, the loss of SIB1 significantly delays RCD (Aviv et al., 2002; Lv et al., 2019), indicating that SIB1 is one of the RCD-triggering components directed by NPR1. The SIB1-GLK interaction in the nucleus enhances the expression of PhANGs, while chloroplast-localized SIB1 (cpSIB1) represses the expression of photosynthesis-associated plastid genes (PhAPGs) (Lv et al., 2019; Morikawa et al., 2002). This concurrent uncoupled expression of PhANGs and PhAPGs increases singlet oxygen (^1^O_2_) levels in chloroplasts through enhanced photoinhibition in PSII (Lv et al., 2019). EXECUTER 1 (EX1), a ^1^O_2_ sensor protein (Dogra et al., 2019), then mediates ^1^O_2_-triggered RS to contribute to stress responses in *lsd1* mutant plants (Lv et al., 2019). It appears that SIB1 undergoes co-translational N-terminal acetylation (NTA) and post-translational ubiquitination (Li et al., 2020). While NTA renders the nuclear SIB1 (nuSIB1) more stable, the latter modification promotes its turnover via the ubiquitin-proteasome system (UPS). The interplay of NTA and UPS seems to regulate nuSIB1-mediated stress responses finely. Nonetheless, earlier reports regarding the positive role of both nuSIB1 and cpSIB1 to RCD suggest that LSD1 may be required to repress the expression of PhANGs to sustain ^1^O_2_ homeostasis.

Here, we demonstrate that LSD1, a transcription coregulator and negative regulator of cell death, interacts with GLK1/2. LSD1 considerably diminishes the GLK binding activity to promoters of the examined PhANGs in Arabidopsis. In agreement, LSD1-overexpressing plants exhibit significantly reduced levels of PhANGs, whereas loss of LSD1 causes a notable upregulation of PhANGs relative to WT plants. SA most likely intervenes in the LSD1-GLK interaction through a rapid accumulation of nuSIB1, leading to a nuSIB1-GLKs interaction, enhanced expression of PhANGs, and activation of EX1-dependent ^1^O_2_ signaling implicated in cell death. We thus concluded that the stress-associated but mutually exclusive transcription coregulators nuSIB1 (positive regulator) and LSD1 (negative regulator) antagonistically regulate the expression of PhANGs through the physical interaction with GLKs. Such antagonistic regulation of GLK activity by nuSIB1 and LSD1 might be instrumental in sustaining ^1^O_2_ homeostasis under SA-associated stress conditions.

## RESULTS

### LSD1 interacts with the GOLDEN2-LIKE transcription factors GLK1 and GLK2

The stress hormone SA primes cell death in the *lsd1* mutant in a light-dependent manner, a typical characteristic of most lesion mimic mutants, as manifested by the abrogated cell death by either loss of key SA signaling components (such as NPR1) or overexpression of the bacterial salicylate hydroxylase NahG that metabolizes SA (Lv et al., 2019; Muhlenbock et al., 2008). Upon exposure to various stimuli, including light, cold, UV-C, red light, hypoxia, and pathogens (Chai et al., 2015; Dietrich et al., 1997; Huang et al., 2010; Jabs et al., 1996; Karpinski et al., 2013; Muhlenbock et al., 2007; Muhlenbock et al., 2008; Rusaczonek et al., 2015), *lsd1* mutant plants drastically develop the foliar RCD phenotype. Among those differentially regulated genes prior to the onset of RCD, the SA-induced transcription coregulator nuSIB1 potentiates the expression of PhANGs and stress-related genes by modulating the TF activity of GLK1/2 and WRKY33, respectively (Lai et al., 2011; Lv et al., 2019; Zarrinpar et al., 2003). These data suggest a possible antagonism between LSD1 and nuSIB1 because nuSIB1-driven stress responses occur in the absence of LSD1. In this regard, we sought if LSD1 also interacts with GLK1/2 to modulate the expression of PhANGs.

We then generated Arabidopsis WT transgenic plants overexpressing GREEN FLUORESCENT PROTEIN (GFP)-tagged LSD1 under the control of the CaMV 35S promoter (35S) (hereafter *oxLSD1*) to unveil putative LSD1-associated proteins. The immunoblot assay detected the LSD1-GFP fusion protein at the predicted molecular mass of approximately 46 kD using an anti-GFP antibody (Supplemental Figure 1). Next, using GFP antibody-conjugated magnetic beads, we co-immunoprecipitated LSD1-GFP and its putative associated proteins from the transgenic plants. The trypsin-digested protein samples were then subjected to tandem mass spectrometry (MS) analyses. The co-immunoprecipitation (Co-IP) coupled to MS analysis using three independent biological replicates identified 217 proteins, which were detected in at least two independent biological replicates, but absent in protein samples of WT and GFP-overexpressing transgenic plants (*35S:GFP*) (Supplemental Dataset 1).

**Figure 1.**
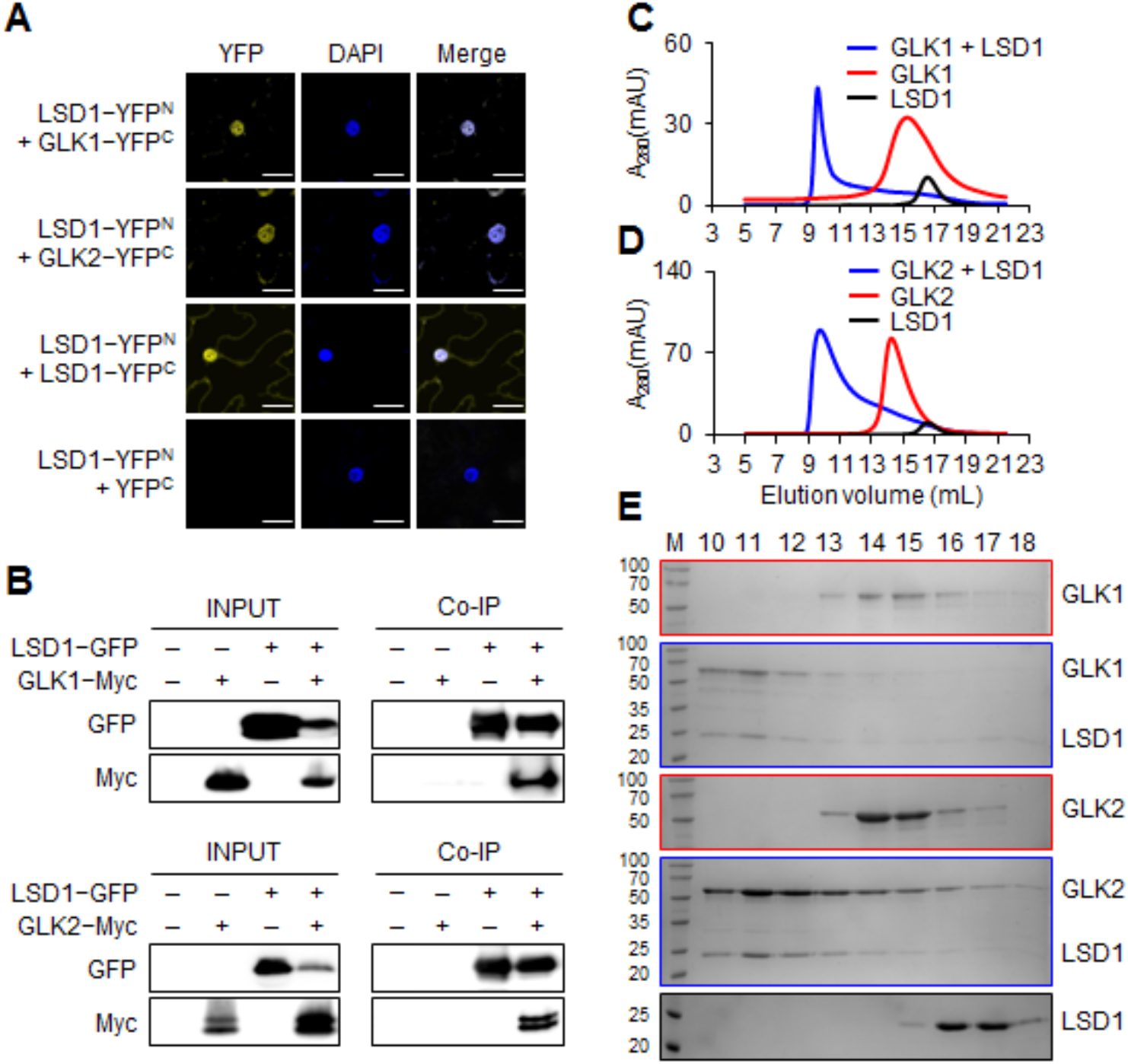
LSD1 interacts with GLK1 and GLK2. **(A)** Bimolecular fluorescence complementation (BiFC) assays. The GLK1 or GLK2 fused with the C-terminal part of YFP (YFP^C^) were coexpressed with the N-terminal part of the YFP (YFP^N^) fused with LSD1 in *N. benthamiana* leaves. The combinations of LSD1-YFP^N^ + LSD1-YFP^C^ and LSD1-YFP^N^ + YFP^C^ were used as a positive and negative control, respectively. DAPI was used to stain the nucleus. All images were taken at the same scale (scale bars: 25 μm). **(B)** Co-immunoprecipitation (Co-IP) analyses using *N. benthamiana* leaves transiently coexpressing LSD1-GFP and GLK1-Myc (or GLK2-Myc). Co-IP was performed using GFP-Trap beads, and the interaction was evaluated by using Myc antibody. **(C–E)** Gel filtration assays showing *in vitro* interaction between LSD1 and GLK proteins expressed in *E. coli*. Gel filtration profiles of LSD1, GLK1, and LSD1-GLK1 complex **(C**) and of LSD1, GLK2, and LSD1-GLK2 complex **(D)**. A_280_(mAU), micro-ultraviolet absorbance at the wavelength of 280 nm. Coomassie blue staining of the peak fractions following SDS-PAGE **(E)**. Numbers on top of SDS-PAGE panels indicate elution volume (mL). M, molecular weight ladder (kD).

Accordingly, among the 217 proteins, we identified both GLK1 and GLK2 (Supplemental Dataset 1). In Arabidopsis, GLK1 and its homolog GLK2 share around 50% amino acid sequence identity. Both contain two conserved domains, a DNA-binding domain (DBD) and a GLK1/2-specific C-terminal GCT-box (Supplemental Figure 2) (Fitter et al., 2002; Rossini et al., 2001). A domain comparison between GLK1 and GLK2 shows a 90% and 79% identity, respectively (Bravo-Garcia et al., 2009). Therefore, it is not surprising that both GLK1 and GLK2 were detected as putative LSD1-associated proteins. Next, we performed a bimolecular fluorescence complementation (BiFC) assay in *Nicotiana benthamiana* (*N. benthamiana*) leaves. Consistent with the previous report (Czarnocka et al., 2017), we confirmed the LSD1-LSD1 interaction in the nucleus, as evident in the overlapped signals detected from YELLOW FLUORESCENT PROTEIN (YFP) and blue-fluorescent DNA stain 4’, 6-diamidino-2-phenylindole (DAPI)-stained nucleus, as well as in the cytosol (Figure 1A). Similarly, we observed a YFP signal in *N. benthamiana* leaf coexpressing LSD1-YFP^N^ and GLK1 (or GLK2)-YFP^C^ (Figure 1A). All YFP signals were exclusively observed in the nucleus (Figure 1A). The Co-IP and an ensuing immunoblot assay further corroborated the LSD1-GLK interaction (Figure 1B). We also purified full-length recombinant proteins of LSD1, GLK1, and GLK2 expressed in *Escherichia coli*. Subsequent gel filtration assays demonstrated that GLK1 (Figure 1C and 1E) and GLK2 (Figure 1D and 1E) form a complex with LSD1, as shown by their co-migration.

### LSD1 interacts with GLK1 and GLK2 through the proline-rich domain

We then generated truncated GLK variants to determine which domain is required for the interaction with LSD1. Prior to the interaction analysis, GLK1/2 and their variants lacking either DBD, potential proline-rich domain (PRD, located between DBD and GCT-box; see discussion), or GCT-box were C-terminally fused with GFP to monitor their nuclear localization (Figure 2A; Supplemental Figure 2). Following transient expression, all intact and variants of Arabidopsis GLK1/2 localized in the nucleus in *N. benthamiana* leaves but with a weak cytosolic GFP signal of GCT-box-deleted GLK1 and GLK2 (Supplemental Figure 3). To examine their interaction with Arabidopsis LSD1 protein, various combinations of BiFC constructs, as shown in Figure 2A, were expressed in *N. benthamiana* leaves to observe their interactions under the confocal microscope. The result clearly showed that the PRD of GLK1/2 is indispensable for the interaction with LSD1 (Figure 2B), further verified by Co-IP analyses (Figure 2C). We then generated GLK1/2 variants by C-terminal serial deletions to ascertain the significance of PRD for the interaction (Supplemental Figure 4A). All GFP-tagged proteins transiently expressed in *N. benthamiana* leaves were localized to the nucleus (Supplemental Figure 4B). The resulting BiFC and Co-IP analyses confirmed the critical role of PRD for LSD1 interaction, as evidenced by the lack of YFP signal when coexpressing LSD1 and GLK1/2 variants lacking the PRD-including C-terminal part (Supplemental Figure 4C and 4D).

**Figure 2.**
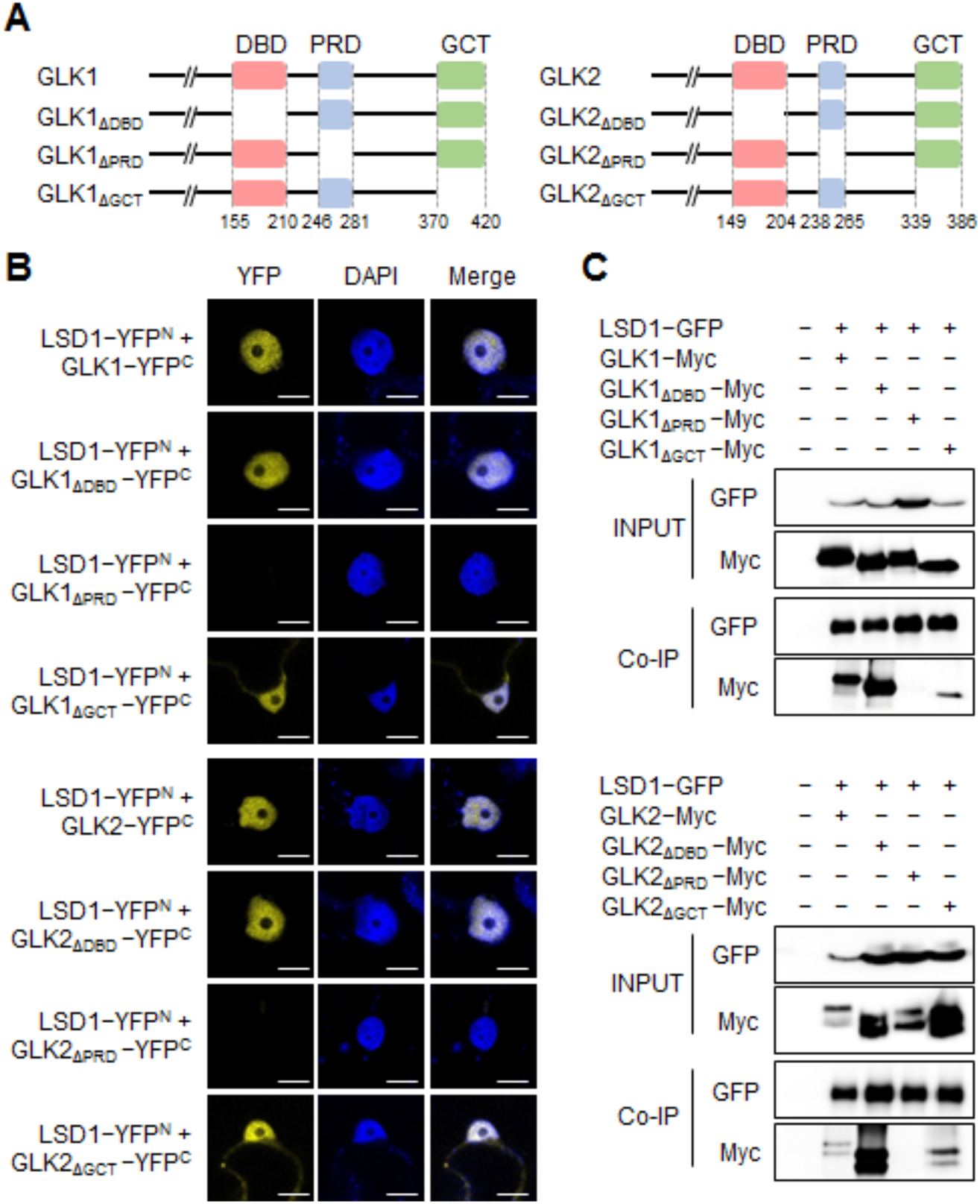
PRD is indispensable for the interaction with LSD1. **(A)** Schematic diagrams show intact GLK1/2 as well as their domain-deleted variants. **(B)** BiFC assays. Each of intact and domain-deleted GLK variants fused with YFP^C^ was coexpressed with LSD1 fused with YFP^N^ in *N. benthamiana* leaves. DAPI was used to stain the nucleus. All images were taken at the same scale (scale bars: 10 μm). **(C)** Co-IP analyses using *N. benthamiana* leaves transiently coexpressing LSD1-GFP with the indicated domain-deleted variant of GLK1/2 fused with Myc-tag.

### Loss of *LSD1* upregulates GLK target genes

Given that GLKs promote the expression of PhANGs (Waters et al., 2009), it is plausible that loss of LSD1 may primarily affect their abundance. To identify affected genes either by loss of LSD1 or of GLK1/2, we compared the RNA sequencing (RNA-seq) data of *lsd1* versus WT (Lv et al., 2019) and *glk1 glk2* versus WT (Ni et al., 2017). As shown in Supplemental Figure 5A, a total of 91 genes (Supplemental Dataset 2) are shared between the upregulated genes (395, at least twofold) in *lsd1* versus WT (Supplemental Dataset 3) and the downregulated genes (936, at least twofold) in *glk1 glk2* versus WT (Supplemental Dataset 4). The Gene Ontology enrichment analysis with the 91 genes for the biological process revealed that photosynthesis and light-harvesting in PSII are over-represented (*P*-value=1.42E-07) (Supplemental Figure 5B; Supplemental Dataset 5). The potentiated expression of PhANGs in *lsd1* versus WT plants is indicative of a negative role of LSD1 in GLK activity. To this end, we also examined the transcript abundance of PhANGs in WT, *glk1 glk2*, and two independent *oxLSD1* transgenic lines (Supplemental Figure 6A) using reverse transcription-quantitative PCR (RT-qPCR). The results indicated that the examined GLK target genes, such as genes encoding light-harvesting chlorophyll *a/b* binding proteins (LHCBs in PSII and LHCA1 in PSI) and chlorophyll synthesis enzymes were substantially repressed in *oxLSD1* relative to WT plants (Figure 3A). *oxLSD1* plants exhibited comparable levels of *GLK1* and *GLK2* transcripts relative to WT (Supplemental Figure 6B), implying that the repression of PhANGs likely resulted from the post-translational regulation of GLK1/2. Remarkably, the overexpression of LSD1-GFP fusion proteins in WT prematurely terminated chloroplast development, which is reminiscent of the phenotype observed in *glk1 glk2* double mutant (Figure 3B and 3C). Some mesophyll cells with nearly undetectable LSD1-GFP signals showed WT-like chloroplasts (Figure 3B). One explanation might be an ectopic cosuppression of the transgene, which also dilutes the molecular phenotypes (e.g., the transcript levels of PhANGs) in the examined leaf tissue. Two independent *oxLSD1* lines with higher *LSD1* transgene expression than the endogenous *LSD1* in WT plants (Figure 3D) exhibited similar phenotypes, such as partial cosuppression of the transgene, defect in chloroplast biogenesis, and reduced levels of LHCB proteins (Figure 3B, 3C, and 3E). Regardless of the promoters used (35S and native), all stable transgenic lines (over 20 lines) showed nearly undetectable or detectable GFP signals but with partial cosuppression (data not shown).

**Figure 3.**
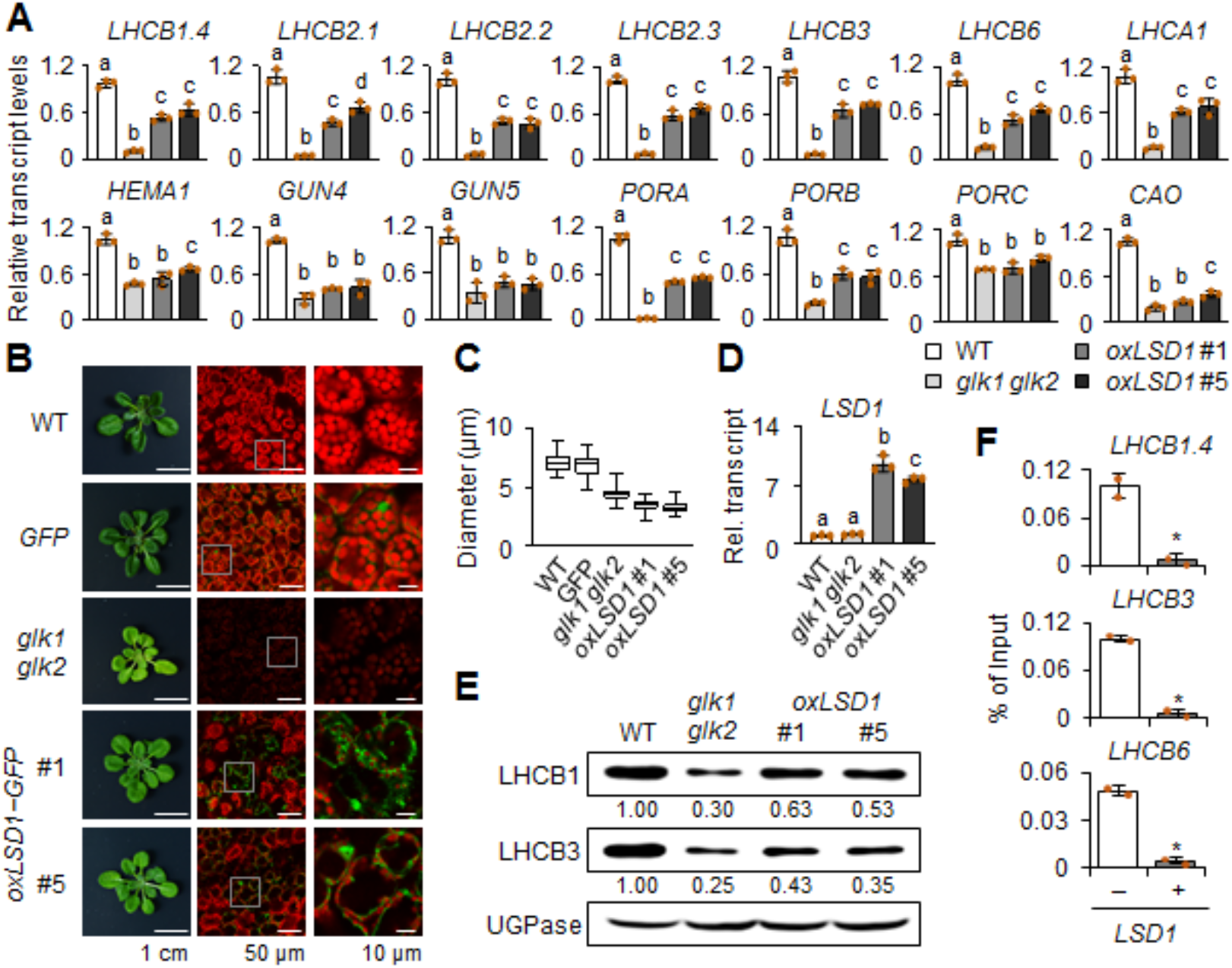
Overexpression of LSD1 negatively affects the expression of GLK target genes and chloroplast biogenesis. **(A)** Relative transcript levels of GLK1 and GLK2 target genes, such as *LHCBs, LHCA1*, and chlorophyll synthesis genes including *glutamyl tRNA reductase (HEMA1/Glu-TR), genome uncoupled 4 (GUN4), magnesium chelatase H subunit (CHLH/GUN5), protochlorophyllide oxidoreductase (POR) A, PORB, PORC*, and *chlorophyllide a oxygenase* (*CAO*) were examined in CL-grown 24-d-old WT, *glk1 glk2*, and *oxLSD1* (#1 and #5) plants using RT-qPCR. **(B)** Plant phenotypes (left panels) and GFP fluorescence (green) of LSD1-GFP fusion proteins merged with chlorophyll autofluorescence signals (red; middle and right panels) in 24-d-old WT, *glk1 glk2*, and transgenic WT plants overexpressing GFP alone (*GFP*) or LSD1-GFP (*oxLSD1* #1 and #5) grown under CL conditions. The small white square boxes in the middle panels were enlarged (right panels). **(C)** Means of chloroplast diameter. At least three confocal images taken from three independent leaves were used to measure the chloroplast diameter. For *oxLSD1*, only mesophyll cells with detectable GFP signals were chosen. **(D and E)** Relative levels of *LSD1* transcript **(D)** and LHCB proteins **(E)** in 24-d-old CL-grown plants of WT, *glk1 glk2*, and *oxLSD1* #1 and #5. UGPase was used as a loading control for the immunoblot analysis in **(E)**. Numbers at the bottom of each immunoblot result indicate the relative quantities of LHCB1 or LHCB3 proteins against the control signal of UGPase. For the RT-qPCR analyses In **(A)** and **(D)**, *ACT2* was used as an internal standard. The value represents means ± standard deviation (SD) (n=3). Lowercase letters indicate statistically significant differences between mean values (P < 0.05, one-way ANOVA with posthoc Tukey’s HSD test). **(E)** ChIP–qPCR results showing the effect of LSD1 overexpression on GLK1 binding to the promoter regions of its target genes (*LHCB1.4, LHCB3*, and *LHCB6*). Myc-tagged GLK1 (GLK1-Myc) was transiently expressed with (+ LSD1) or without LSD1-RFP (− LSD1) in Arabidopsis leaf protoplasts isolated from *lsd1 glk1 glk2* triple mutant. The enrichment value was normalized to the input sample, representing means ± SD from two independent ChIP assays. Asterisks denote statistically significant differences by Student’s *t-*test (*P* < 0.01) from the value of - LSD1.

### LSD1 inhibits the DNA-binding activity of GLK1

Regarding that nuclear-localized LSD1 acts as a transcription coregulator and that LSD1 interacts with GLKs, it is conceivable that LSD1 might directly regulate the DNA-binding activity of GLK1/2. Therefore, we performed a chromatin immunoprecipitation (ChIP) coupled with a qPCR analysis. The relative activity of GLK1 towards its target promoters was examined in the presence or absence of LSD1 using protoplasts isolated from the rosette leaves of *lsd1 glk1 glk2* triple mutant plants. Since GLK1 and GLK2 are highly unstable (Tokumaru et al., 2017; Waters et al., 2008), we used a protoplast transient expression system to ensure sufficient protein expression to elucidate the impact of LSD1 on GLK1 function. The *35S:GLK1-Myc* was transiently coexpressed with either *35S:RED FLUORESCENT PROTEIN* (*RFP*) or *35S:LSD1-RFP* in the protoplasts. ChIP assays were then performed with nuclear lysis from transfected protoplasts. With anti-Myc antibody-conjugated agarose beads, the Myc-tagged protein-DNA complex was pulled down. The immunoprecipitated DNA was then analyzed using qPCR to compare the DNA-binding activity of GLK1 in the presence or absence of LSD1. Afterward, we examined the relative expression levels of well-established GLK target genes such as *LHCB1.4, LHCB3,* and *LHCB6* (Waters et al., 2009). The results demonstrated that the presence of LSD1 markedly diminished the DNA-binding activity of GLK1 to promoters of these *LHCB* genes (Figure 3F).

### Loss of LSD1 potentiates ^1^O_2_-triggered EX1-dependent RS in *flu* mutant

We next validated the above ChIP assay result *in planta*. Considering the positive regulation of chlorophyll synthesis by GLK1 (Waters et al., 2009) and the repression of GLK activity by LSD1 (Figure 3A and 3F), it is tempting to hypothesize that loss of LSD1 might increase the rate of chlorophyll biosynthesis. By revisiting the previously published RNA-seq data (Lv et al., 2019), we noticed that a set of chlorophyll synthesis genes including *glutamyl tRNA reductase* (*HEMA1/Glu-TR*), *genome uncoupled 4* (*GUN4*), *magnesium chelatase H subunit* (*GUN5/CHLH*), *magnesium protoporphyrin IX monomethyl ester cyclase* (*CHL27/CRD1*), *protochlorophyllide oxidoreductase A* (*PORA*), and *chlorophyllide a oxygenase* (*CAO*) were markedly upregulated in *lsd1* before the onset of RCD (Figure 4A and 4B). We then analyzed the 5-aminolevulinic acid (5-ALA, the common precursor of all tetrapyrroles) synthesis rate in light-grown *lsd1* mutant plants treated with levulinic acid (LA) (Nandi and Shemin, 1968), a competitive chemical inhibitor of 5-ALA dehydratase that catalyzes the synthesis of porphobilinogen through the asymmetric condensation of two 5-ALA molecules (Figure 4A). It is important to note that the 5-ALA synthesis is the rate-limiting step for chlorophyll synthesis (Beale and Castelfranco, 1974; Hou et al., 2019) (Figure 4A). In view of the fact that FLUORESCENT (FLU) protein directly represses Glu-TR activity to inhibit protochlorophyllide (Pchlide) accumulation in the dark (Goslings et al., 2004) (Figure 4A) and that *flu* mutant plants exhibit a higher 5-ALA synthesis rate in the presence of LA under continuous light (CL) conditions (Goslings et al., 2004), we used *flu* as a positive control.

**Figure 4.**
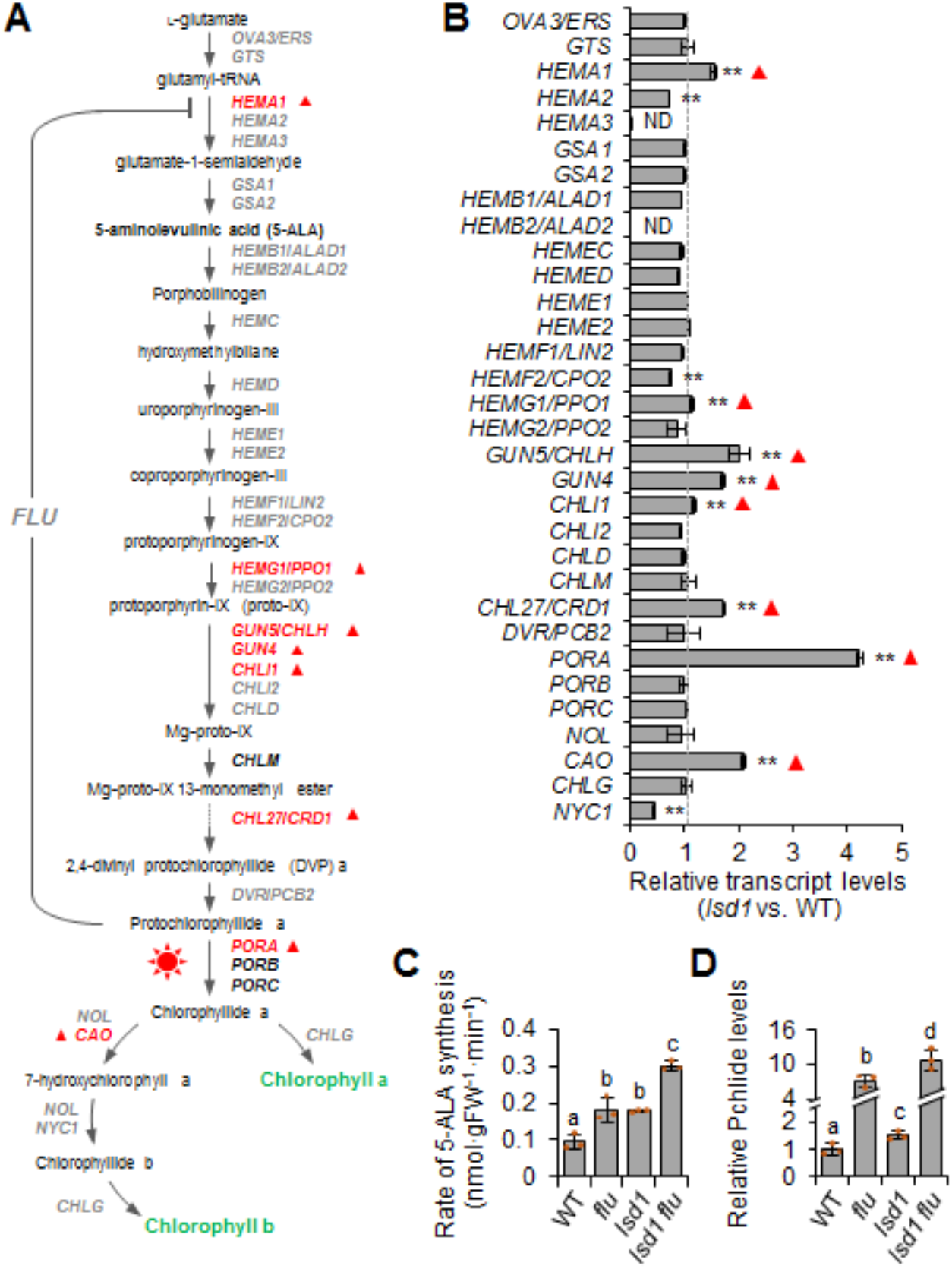
*LSD1* mutation leads to the transcriptional upregulation of chlorophyll synthesis genes. **(A)** A schematic representation of the chlorophyll synthesis pathway. **(B)** The expression levels of chlorophyll synthesis genes represented in **(A)** were obtained from our previous study (Lv et al., 2019), and the relative transcript levels of chlorophyll synthesis genes in 17-d-old CL-grown *lsd1* mutants compared to wild type (WT) are represented. Error bars indicate SD (n=3). Asterisks indicate statistically significant differences (\**P* < 0.05; \*\**P* < 0.01) in the *lsd1* mutant determined by Student’s *t*-test relative to wild type. ND: non-detected. Red triangles in **(A)** and **(B)** indicate the significantly upregulated genes in the *lsd1* mutant compared to WT. **(C)** Levels of 5-ALA synthesis rate under CL conditions. The 5-ALA synthesis rate was measured in 16-d-old plants of WT, *flu, lsd1*, and *lsd1 flu*. **(D)** Relative levels of protochlorophyllide (Pchlide) in 10-d-old plants of WT, *flu, lsd1*, and *lsd1 flu* grown under CL and then transferred to the dark for 8 hours. Values in **(C)** and **(D)** represent means ± SD (n=3). Lowercase letters indicate significant differences between the indicated genotypes (*P* < 0.05, one-way ANOVA with posthoc Tukey’s HSD test).

The 5-ALA synthesis rate was almost comparable in *lsd1* and *flu* seedlings in the presence of LA (Figure 4C). Although it is yet unclear whether the transcriptional upregulation of *HEMA1* is responsible for the 5-ALA accumulation in *lsd1*, the concurrent loss of both *FLU* and *LSD1* further increased the 5-ALA synthesis rate under CL conditions (Figure 4C). Since Pchlide levels in the dark would indirectly reflect chlorophyll biosynthesis rate owing to the absence of the enzyme(s) involved in Pchlide turnover (Forreiter and Apel, 1993), we measured Pchlide levels in dark-incubated plants of WT, *flu, lsd1*, and *lsd1 flu* using high-performance liquid chromatography analysis. As anticipated, Pchlide was highly upregulated in the *flu* mutant background, as demonstrated earlier (Meskauskiene et al., 2001) (Figure 4D). The loss of LSD1 raises the Pchlide level (approximately a 1.5-fold increase) in both WT and *flu* mutant backgrounds. The presence of FLU protein in *lsd1* seems to prevent the drastic accumulation of Pchlide in the dark. Collectively, these results corroborate the negative role of LSD1 towards GLK activity, which seems to be, at least in part, required for tetrapyrrole homeostasis.

We then hypothesized that the SA-induced nuSIB1-GLK-driven upregulation of PhANGs and the higher chlorophyll synthesis rate by *FLU* mutation might further enhance ^1^O_2_ levels in chloroplasts in *lsd1 flu* plants grown under CL conditions before the onset of RCD. Indeed, *lsd1 flu* double mutant plants exhibited accelerated RCD than *lsd1* plants (Figure 5A), which was found to be radically reduced in *lsd1 flu ex1*, indicating ^1^O_2_ was the prime cause of the reinforced RCD in *lsd1 flu*. This result coincided with the intensity of maximum fluorescence of PSII (Fm) (Figure 5A), PSII maximum efficiency (Fv/Fm) (Figure 5B), and the abundance of ^1^O_2_-responsive genes (SORGs) (Dogra et al., 2017) (Figure 5C).

**Figure 5.**
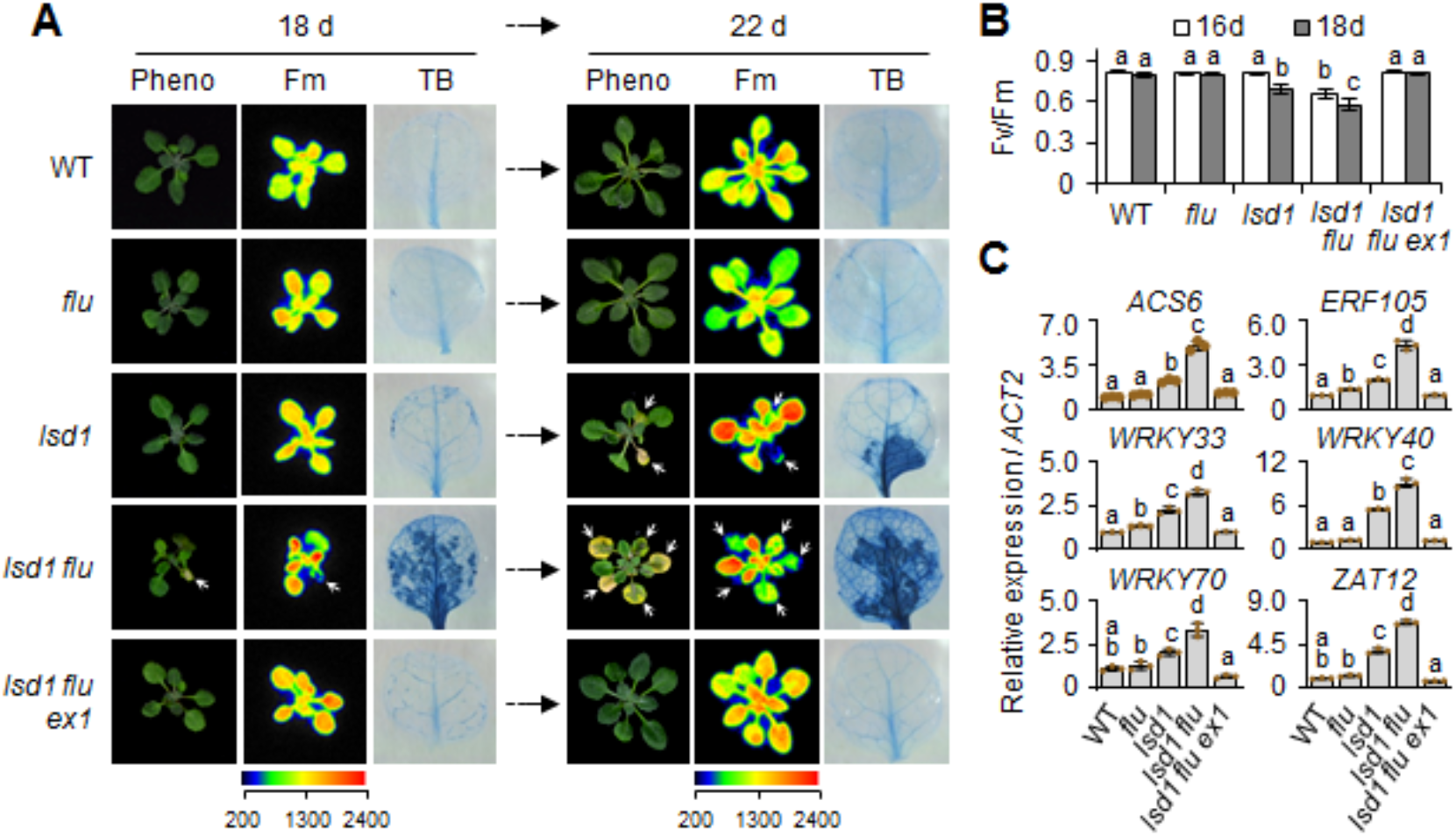
Loss of LSD1 potentiates ^1^O_2_-triggered EX1-dependent cell death in *flu* mutant. **(A)** WT, *flu, lsd1, lsd1 flu*, and *lsd1 flu ex1* plants were grown on Murashige and Skoog (MS) medium under CL conditions (100 μmol·m^-2^·s^-1^). The RCD phenotype (Pheno, left panels) and the chlorophyll maximum fluorescence (Fm) of PSII (middle panel) were monitored in the whole plants at the indicated time points. The dead cells in the first or second leaves from the genotypes were visualized via trypan blue staining (right panel). Images are representative phenotypes. **(B)** The first or second leaves from each genotype were harvested at the indicated time points to measure the maximum photochemical efficiency of PSII (Fv/Fm). Data represent means ± SD (n=10). **(C)** Expression levels of selected ^1^O_2_-responsive genes (SORGs) were examined by RT-qPCR in 18-d-old plants. *ACT2* was used as an internal standard. Data represent means ± SD (n=3). Lowercase letters in (**B**) and **(C)** indicate statistically significant differences between mean values (P < 0.05, one-way ANOVA with post-hoc Tukey’s HSD test).

### SA-induced SIB1 interrupts LSD1-GLK1 interaction

Since SIB1-mediated genomes uncoupled expression of PhANGs and PhAPGs largely contributes to *lsd1* RCD via ^1^O_2_ signaling (Lv et al., 2019), we hypothesized that nuSIB1 would counteractively modulate LSD1-GLK interaction to reinforce the expression of PhANGs. Considering its rapid turnover via UPS (Li et al., 2020), nuSIB1 may promptly intervene in this LSD1-GLK interaction, resulting in nuSIB1-GLK interaction and reinforced expression of PhANGs, thereby contributing to cell death (Lv et al., 2019). Alternatively, SA per se may interfere with LSD1-GLK interaction, for instance, through alteration of protein conformation of LSD1 or GLKs or both. In fact, a previous report showed a redox-sensitive reconfiguration of LSD1 and concurrent change of its interactome (Czarnocka et al., 2017). Thus, we examined how SA impacts the LSD1-GLK1 interaction in Arabidopsis leaf protoplasts isolated from WT and *sib1* mutant plants. The result that SA significantly hindered the LSD1-GLK1 interaction in WT but not in *sib1* (Figure 6A) suggested that nuSIB1 rather than SA per se interrupts LSD1-GLK1 interaction. To further elucidate an antagonistic action of nuSIB1 towards LSD1-GLK1 interaction, a dose-dependent impact of nuSIB1 was examined. For this, LSD1-GFP and GLK1-Myc were transiently coexpressed in Arabidopsis leaf protoplasts, along with different amounts of free RFP or SIB1-RFP. It should be noted that increasing doses of RFP or SIB1-RFP reduce the expression of LSD1-GFP and GLK1-Myc, probably as a consequence of diminished transfection efficiency due to the presence of the additional constructs (Figure 6B). Nonetheless, the relative amount of GLK1-Myc protein co-immunoprecipitated with LSD1-GFP was quantified using ImageJ following immunoblot analysis (Figure 6C). The results showed a SIB1 dose-dependent inhibition of the LSD1-GLK1 interaction.

**Figure 6.**
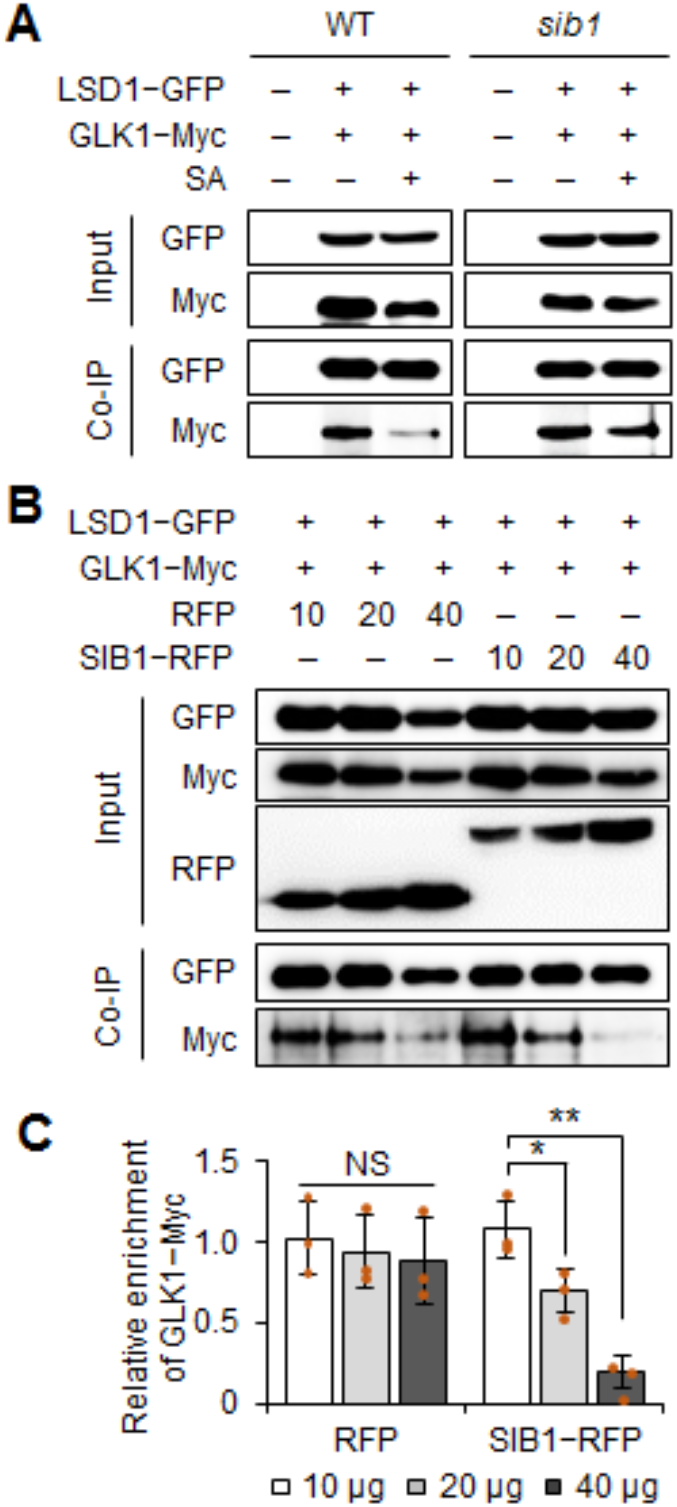
SA-induced SIB1 intervenes in LSD1-GLK interaction. **(A)** The effect of the SA-induced nuSIB1 on the LSD1-GLK1 interaction. For Co-IP analyses, *35S:LSD1-GFP* and *35S:GLK1-Myc* were transiently coexpressed in Arabidopsis leaf protoplasts. The protoplasts were treated with either mock or 0.2 mM SA for 5 hours. **(B)** The dose-dependent impact of SIB1 on the LSD1-GLK1 interaction. As indicated, *35S:LSD1-GFP* and *35S:GLK1-Myc* were coexpressed in Arabidopsis leaf protoplasts isolated from WT plants together with different amounts (10 μg, 20 μg, or 40 μg, respectively) of a plasmid containing either free *35S:RFP* or *35S:SIB1-RFP*. The subsequent Co-IP and immunoblot results are shown. Three independent experiments were conducted with similar results, and representative results are shown in **(A)** and **(B)**. **(C)** The signal intensity of eluted GLK1-Myc (from triplicate immunoblots in **B**) versus its input signal was quantified using the ImageJ software. Data are means ± SD (n=3). Asterisks denote statistically significant differences by Student’s *t*-test (\**P* < 0.05, \*\**P* < 0.01, NS: not significant).

One possible scenario for the nuSIB1-dependent interruption of LSD1-GLK interaction is that SA-induced nuSIB1 may directly interact with LSD1, releasing GLK1 and GLK2 in the nucleus. The free GLK1/2 may interact with excess nuSIB1, promoting the expression of PhANGs. However, while the LSD1-LSD1 interaction was apparent, no interaction between LSD1 and nuSIB1 was observed (Supplemental Figure 7). Then we assumed that SA-induced nuSIB1 might compete with LSD1 to bind to the PRD of GLK1 and GLK2. We then carried out Co-IP analyses to investigate if PRD is required for the interaction with SIB1. The result showed that the N-terminal region excluding all three domains is sufficient to interact with nuSIB1 (Supplemental Figure 8A and 8B). Since the N-terminal part contains a nuclear localization signal (Zhang et al., 2021a), we ended further defining the minimum length of the N-terminal necessitated for the interaction with nuSIB1. It is likely that SA-induced nuSIB1 competitively interacts with GLK1 and GLK2 through the N-terminal part, which consequently enhances the expression of PhANGs and the ^1^O_2_ level, thereby activating an EX1-mediated cell death response (Figure 7). The rapid turnover of nuSIB1 via UPS (Li et al., 2020) might result in LSD1-GLK interaction and restore the expression levels of PhANGs.

**Figure 7.**
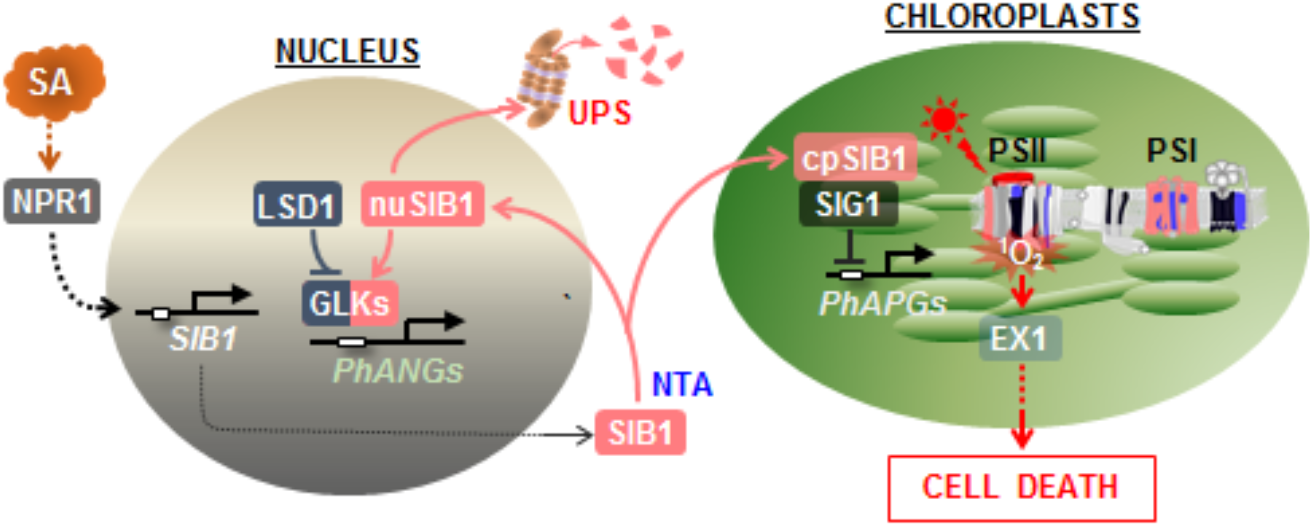
Proposed model elucidating the counteractive regulation of GLKs by LSD1 and nuSIB1. LSD1-GLK interaction is required for negative regulation of GLK activity to fine-tune the expression of PhANGs, including LHCBs and chlorophyll synthesis genes. Under SA-increasing stress conditions, the NPR1-induced and NTA-stabilized nuSIB1 intervenes in LSD1-GLK interaction to reinforce the expression of PhANGs, while the cpSIB1 represses the expression of PhAPGs by interacting with SIG1 (Li et al., 2020; Lv et al., 2019; Morikawa et al., 2002). The resulting uncoupled expression of PhANGs and PhAPGs aggravates PSII photoinhibition and increases ^1^O_2_ level in chloroplasts, enabling EX1-mediated retrograde signaling to activate the expression of SORGs and cell death response (Kim et al., 2012; Lv et al., 2019). While NTA renders nuSIB1 more stable, UPS promotes the proteolysis of nuSIB1 (Li et al., 2020), restoring LSD1-GLK interaction to avoid an excess of PhANG expression and ^1^O_2_ accumulation. The counteractive regulation of GLKs by nuSIB1 and LSD1, along with post-translational regulation of nuSIB1 stability, seems vital to modulate ^1^O_2_ levels in chloroplasts during and after SA-increasing stress conditions.

## DISCUSSION

Besides their essential role in chloroplast biogenesis and photosynthesis, multiple lines of evidence demonstrate that GLKs function in plant stress responses, evoking an intriguing proposal that GLK may serve as a master switch in synchronously regulating photosynthesis and stress responses. We previously reported that the positive regulator of SA signaling and transcription coregulator nuSIB1 interacts with GLKs and WRKY33 to reinforce the expression of PhANGs and SA-responsive genes, respectively, upon an increase in cellular SA level (Li et al., 2020; Lv et al., 2019). On the contrary, cpSIB1 interacts with SIG1 polymerase to repress the expression of PhAPGs (Lv et al., 2019; Morikawa et al., 2002; Xie et al., 2010). The genomes-uncoupled expression of PhANGs and PhAPGs heightens the PSII photoinhibition, thereby escalating the highly reactive oxygen species, specifically ^1^O_2_. ^1^O_2_ then contributes to SA-driven plant stress responses via EX1-mediated RS, which is shown to reinforce RCD phenotype in *lsd1* mutant (Dogra et al., 2019; Lv et al., 2019). Since the SA receptor NPR1 is required to induce the expression of *SIB1* (Xie et al., 2010), EX1-mediated ^1^O_2_ signaling is likely to be one of the downstream events led by SA and NPR1.

We now showed that LSD1 interacts with GLK1 and GLK2 TFs in the nucleus (Figure 1A). Besides their transcriptional regulation (e.g., by GUN1-mediated RS), multiple proteins post-translationally modulate GLK activity (Tang et al., 2016; Tokumaru et al., 2017; Zhang et al., 2021a). The C-terminal GCT-box drives GLK homo-or hetero-dimerization in maize (Rossini et al., 2001). The *turnip yellow mosaic virus* (TYMV) protein P69 binds to the GLK1/2 GCT-box, repressing PhANGs and chloroplast biogenesis in Arabidopsis (Ni et al., 2017). On the contrary, BRASSINOSTEROID INSENSITIVE 2 (BIN2)-dependent GLK phosphorylation promotes chloroplast biogenesis by stabilizing GLK proteins in Arabidopsis (Zhang et al., 2021a). These reports indicate that both DBD and GCT-box in GLK1/2 are involved in protein-protein interaction. Notably, the interdomain region of GLK1 and GLK2 are proline-enriched (Figure 2A; Supplemental Figure 2). Since proline residues provide protein-docking sites (Siligardi and Drake, 1995; Zarrinpar et al., 2003), we anticipated the PRD as an additional candidate domain required for the interaction with LSD1. The ensuing BiFC and Co-IP assays verified that the PRD is central for interacting with LSD1. GLK1 and GLK2 lacking DBD and GCT-box but retaining PRD interacted with LSD1, but complete loss of PRD abolished these interactions (Figure 2B and 2C). Besides, the association of GLK2 with the CUL4-DDB1-based E3 ligase complex promotes UPS-mediated GLK2 turnover in tomato (Tang et al., 2016). The COP1 and UPS-mediated GLK1 degradation was also reported in Arabidopsis plants with long-term abscisic acid (ABA) treatment (Lee et al., 2021). Interestingly, one latest work showed that WRKY75 directly represses *GLK* expression during leaf senescence (Zhang et al., 2021b). The ABA-induced SIB1 and its close homolog SIB2 interact with and inhibit WRKY75 activity, enabling the expression of GLKs in response to ABA. The antagonistic regulation of GLKs expression by SIB1/2 and WRKY75 was proposed to be essential in controlling ABA-mediated leaf senescence and seed germination. These findings by other groups and our data suggest that the GLKs are common targets of development or stress signaling to modulate chloroplast homeostasis. As emerging notion strongly supports the role of chloroplasts as environmental sensors, such modulation of GLK activity and stability would also significantly affect chloroplast-mediated plant stress responses.

Loss of LSD1 potentiated the expression of GLK target genes such as PhANGs (Supplemental Figure 5A and 5B) and increased the 5-ALA synthesis rate compared to WT plants (Figure 4C). Conversely, LSD1 overexpression repressed the expression of PhANGs (Figure 3A). These results were consistent with *oxLSD1* plant phenotypes exhibiting prematurely terminated chloroplast development and reduced LHCB levels (Figure 3B–3E). Consistently, LSD1 overexpression repressed GLK1 binding activity to its target promoters (Figure 3F). The effects of loss- and gain-of-function of LSD1 towards the expression of GLK target genes also suggest a steady-state LSD1-GLK interaction in WT plants grown under normal growth conditions. Given that the elevated expression of PhANGs directed by nuSIB1-GLK interaction contributes to *lsd1* RCD (Lv et al., 2019), it was tempting to hypothesize that SA-induced nuSIB1 interferes with LSD1-GLK interaction. Indeed, the Co-IP assay confirmed the negative impact of nuSIB1 accumulation on LSD1-GLK interaction (Figure 6). It has been shown that the PRD domain provides the sequence-specific docking site for interacting proteins without the requirement of a high-affinity interaction (Saraste and Musacchio, 1994; Siligardi and Drake, 1995; Zarrinpar et al., 2003). The sequence-specific but low-affinity interaction at the proline-rich region might allow a highly reversible interaction between LSD1-GLKs, enabling SA-induced nuSIB1 to rapidly intervene in this interaction through the N-terminus of GLKs (Supplemental Figure 8). Such versatile regulation of nuSIB1 stability and an antagonistic mode of action of nuSIB1 and LSD1 towards GLK1/2 might be vital to maintain ^1^O_2_ homeostasis in chloroplasts and to induce SA-driven stress responses under fluctuating environmental conditions (Figure 7).

Our findings also raise a plausible idea that GUN1-mediated RS would largely contribute to plant stress responses because the signaling primarily represses the expression of *GLK1* and *GLK2* once the foliar plastid function is interrupted (see introduction). Consistently, *gun1* mutant plants exhibit an increased susceptibility towards heat, water, drought, cold, and high-light stresses with enhanced cellular ROS levels (Cheng et al., 2011; Miller et al., 2007; Tang et al., 2014; Zhang et al., 2013; Zhang et al., 2011). The multifaceted interactions between GLK1/2 and the antagonistic modules nuSIB1 and LSD1, as well as other stress-related proteins, may be accountable for the altered *gun1* phenotype to various stress factors. The enhanced expression of *GLK1* and *GLK2* may increase the level of ^1^O_2_ if nuSIB1 is accumulated and intervene in LSD1-GLK interaction in *gun1*. The ^1^O_2_-triggered EX1-mediated RS may then modulate plant stress responses in *gun1* mutant plants under SA-increasing stress conditions. In this regard, a new study of *gun1* may provide further insight into how chloroplast RS pathways mediated by GUN1 and EX1 coordinate SA-mediated plant stress responses through GLK1/2 and ^1^O_2_, respectively.

## Methods

### Plant materials and growth conditions

The seeds used in this study were derived from *Arabidopsis thaliana* Columbia-0 (Col-0) ecotype and were harvested from plants grown under continuous light (CL; 100 μmol·m^-2^·s^-1^) at 22 ± 2 °C. Arabidopsis mutant seeds used in this study, including *lsd1-2* (SALK_042687) (Lv et al., 2019), *glk1 glk2* (*Atglk1.1; Atglk2.1*) (Fitter et al., 2002), and *ex1* (SALK_002088) (Lee et al., 2007) were obtained from the Nottingham Arabidopsis Stock Centre (NASC). *flu5c* has been described previously (Meskauskiene et al., 2001). The double and triple mutants in the *lsd1-2* background including *lsd1 flu, lsd1 flu ex1*, and *lsd1 glk1 glk2* were generated by crossing the homozygous plants. The genotypes of all mutants were confirmed by PCR-based analyses. Primer sequences used for PCR are listed in Supplemental Table 1.

Seeds were surface sterilized with 70% (v/v) ethanol containing 0.05% (v/v) Triton X-100 (Sigma-Aldrich) for 10 min and washed five times with sterile distilled water. The sterile seeds were plated on Murashige and Skoog (MS) medium (Duchefa Biochemie) with 0.7% (w/v) agar (Duchefa Biochemie) and stratified at 4 °C in darkness for two days prior to placing in a growth chamber (CU-41L4; Percival Scientific) with CL condition.

### Generation of *LSD1* overexpression lines

The stop-codon-less *LSD1* coding sequence (CDS) was cloned into the modified pCAMBIA3300 binary vector containing the 35S promoter, a NcoI restriction site, and the *EGFP*. Arabidopsis stable transgenic lines were generated by a floral dip transformation procedure (Clough and Bent, 1998) with *Agrobacterium tumefaciens* strain GV3101. Homozygous transgenic lines were selected on MS medium containing 12.5 mg/L glufosinate-ammonium (Sigma-Aldrich).

### RNA extraction and RT-quantitative PCR (RT-qPCR)

Total RNA was isolated from leaf tissues using the Spectrum Plant Total RNA Kit (Sigma-Aldrich) according to the manufacturer’s instructions. The concentration of RNA was determined using the ultraviolet-visible spectrophotometer (NanoDrop™, Thermo Fisher Scientific), and the quality of RNA was evaluated by measuring the A260/A280 ratio. cDNA synthesis was performed with 1 μg of total RNA using the PrimeScript™ RT Reagent Kit (Takara) following the manufacturer’s instructions. The RT-qPCR was performed on QuantStudio™ Flex Real-Time PCR System (Applied Biosystems) using iTaq Universal SYBR Green PCR master mix (Bio-Rad). The relative transcript level was calculated by the ddCt method (Livak and Schmittgen, 2001) and normalized to the *ACTIN2* (AT3G18780) gene transcript level. The sequences of the primers used for RT-qPCR are listed in Supplemental Table 1.

### Co-immunoprecipitation (Co-IP) assay

Co-IP assays were performed using *Nicotiana benthamiana* or Arabidopsis leaf protoplasts transiently coexpressed with the indicated combination of proteins. For the Co-IP assays in *N. benthamiana*, the *35S:LSD1-sGFP*, *35S:SIB1-sGFP*, *35S:GLK1-4×Myc*, and *35S:GLK2-4×Myc* constructs were created as described previously (Lv et al., 2019). Briefly, pDONR221/Zeo entry vector (Thermo Scientific) containing the stop codon-less full-length CDS of *LSD1, SIB1, GLK1*, or *GLK2* was recombined into the destination vector pGWB605 for C-terminal fusion with sGFP or into pGWB617 for C-terminal fusion with 4×Myc through the Gateway LR reaction (Thermo Scientific). For the *35S:GLK1-4×Myc* (or *sGFP*) and *35S:GLK2-4×Myc* (or *sGFP*) constructs, a linker DNA encoding Gly-Gly-Ser-Gly-Gly-Ser was added between *4xMyc* (or *sGFP)* tag and GLK1 or GLK2 to increase conformational flexibility of the fusion protein as described previously (Tokumaru et al., 2017). The same procedures were used to create the constructs containing CDSs encoding domain-deleted or C-terminally truncated variants of GLK1 and GLK2. The different combinations of selected vectors were coexpressed in 4-week-old *Nicotiana benthamiana* leaves by Agrobacterium-mediated leaf infiltration as previously described by Boruc et al. (2010). For the Co-IP assays in Arabidopsis leaf protoplasts, the *35S:LSD1-sGFP, 35S:GLK1-4×Myc, 35S:SIB1-RFP*, and *35S:LSD1-RFP* were cloned into the pSAT6 vector (Tzfira et al., 2005). The isolation and transfection of Arabidopsis leaf protoplasts were performed as described previously (Yoo et al., 2007). The indicated combination of vectors was cotransfected into protoplasts (3 × 10^6^) isolated from 4-week-old plants of WT or *sib1*.

Total protein was extracted using an IP buffer containing 50 mM Tris-HCl (pH 7.5), 150 mM NaCl, 0.5mM EDTA, 10% (v/v) glycerol, 1% (v/v) Nonidet P-40 (NP-40), 1% deoxycholate, 0.1% (w/v) SDS, 1 × cOmplete protease inhibitor cocktail (Roche), 1 mM PMSF, and 50 μM MG132. The protein extracts were incubated with 20 μL of GFP-Trap magnetic agarose beads (GFP-TrapMA, Chromotek) for 2 h at 4 °C by vertical rotation (10 rpm). After incubation, the beads were washed five times with the washing buffer containing 10 mM Tris-HCl (pH 7.5), 150 mM NaCl, 0.5 mM EDTA, 1 mM PMSF, 50 μM MG132, and 1 × cOmplete protease inhibitor cocktail. The immunoprecipitated proteins were then eluted with 2 × SDS protein sample buffer [120 mM Tris-HCl (pH 6.8), 20% (v/v) glycerol, 4% (v/w) SDS, 0.04% (v/w) bromophenol blue, and 10% (v/v) β-mercaptoethanol] for 10 min at 95 °C. The eluates were subjected to 10% SDS-PAGE gels, and the interaction between coexpressed proteins was examined by immunoblot analyses using a mouse anti-Myc monoclonal antibody (1:10,000; Cell Signaling Technology), a rat anti-RFP monoclonal antibody (1:10,000; Chromotek), and a mouse anti-GFP monoclonal antibody (1:5,000; Roche).

### Protein Extraction and immunoblot analysis

Total proteins were extracted from 100 mg of foliar tissues with the IP buffer and quantified with a Pierce BCA protein assay kit (Thermo Fisher Scientific). Afterward, 20 μg total protein was separated on 10% SDS-PAGE gels and blotted onto Immun-Blot PVDF membrane (Bio-Rad). LSD1-GFP, LHCB1, and LHCB3 were immunochemically detected with mouse anti-GFP (1:10,000; Roche), rabbit anti-LHCB1 (1:5,000; Agrisera), and rabbit anti-LHCB3 (1:5,000; Agrisera) antibodies, respectively. The UDP-glucose pyrophosphorylase (UGPase) detected with rabbit anti-UGPase (1:3,000; Agrisera) was used as a loading control.

### Confocal laser-scanning microscopy

The GFP, YFP, chlorophyll, and 4’, 6’-diamidino-2-phenylindole (DAPI) fluorescence signals were detected by confocal laser-scanning microscopy analysis using TCS SP8 (Leica Microsystems). All the images were obtained and processed with Leica LAS AF Lite software, version 2.6.3 (Leica Microsystems).

### Bimolecular fluorescence complementation (BiFC) assay

BiFC assays were conducted with a split-YFP system in *N. benthamiana* leaves, as described previously (Lee et al., 2020; Lu et al., 2010). Briefly, the pDONR/Zeo entry vectors (Thermo Fisher Scientific) containing CDSs lacking the termination codon of intact forms, domain-deleted, or C-terminally truncated variants of GLK1 and GLK2 were recombined into the destination vector pGTQL1221 through Gateway LR reaction. The same procedure was done to recombine the pDONR221/ZEO entry vector containing the *LSD1* CDS lacking the terminal codon into the pGTQL1211. For the BiFC assay, *A. tumefaciens* mixtures carrying the appropriate constructs were infiltrated into 4-week-old *N. benthamiana* leaves. The presence of YFP fluorescence signals was evaluated by confocal laser-scanning microscopy analysis.

### ChIP-qPCR assays

ChIP assays were performed using Arabidopsis leaf protoplasts as described previously (Lee et al., 2017; Lv et al., 2019; Yoo et al., 2007). Briefly, 1 mg of pSAT6 vectors containing *35S:GLK1-4×Myc* DNA were transfected with or without pSAT6 vector containing *35S:LSD1-RFP* DNA into Arabidopsis leaf protoplasts (2 × 10^7^) isolated from 4-week-old *lsd1 glk1 glk2* triple mutant plants grown under 10-h light/14-h dark conditions at a light intensity of 100 μmol m^-2^s^-1^. Afterward, the protoplasts were incubated at 24 °C for 16 h under dim light conditions. The protoplast chromatins were crosslinked by 1% (v/v) formaldehyde in 1 × PBS (pH 7.4) for 10 min and quenched with 0.1 M glycine for 5 min. After isolating nuclei from the protoplasts, the chromatins were sheared by sonication into an average size of around 500 bp. The lysates were diluted with 10 × ChIP dilution buffer [1% (v/v) Triton X-100, 2 mM EDTA, 20 mM Tris-HCl (pH 8.0), 150 mM NaCl, 50 μM MG132, 1 mM PMSF, and 1 × protease inhibitor cocktail] and precleared by incubation with 50 μL Protein-A agarose beads/Salmon sperm DNA (Millipore) at 4 °C for 1 h. The samples were then incubated with anti-Myc monoclonal antibodies (1:10,000; Cell Signaling Technology) at 4 °C overnight. To determine non-specific binding of DNA on beads, ChIP assays were also performed without antibodies. After washing the beads, the immunocomplexes were eluted with elution buffer containing 1% (w/v) SDS and 100 mM NaHCO_3_. The eluates were treated with proteinase K for 1 h at 37 °C after reverse cross-linking. The bound DNA fragments were purified as previously described by Lee et al. (2017) and precipitated with ethanol in the presence of glycogen. The purified DNA was dissolved in water. qPCR analyses were performed on bound and input DNAs. The primers for each tested gene are listed in Supplemental Table 1. The amount of DNA enriched by the anti-Myc antibody was calculated in comparison with the respective input DNA used for each ChIP analysis. Afterward, the enrichment was calculated by normalizing against the corresponding control sample (without antibody).

### Production of recombinant proteins

To produce recombinant proteins of LSD1, GLK1, and GLK2, the coding sequences (CDS) of genes were cloned into the modified pET21b (Novagen) expression vector after adding a cleavage site for the tobacco etch virus (TEV) protease to the 5’ end of the CDSs. The recombinant proteins with a cleavable N-terminal 10×His-MsyB tag were expressed in *E. coli* BL21 (DE3). After culturing the cells at 37 °C until an OD_600_ of 0.6, recombinant proteins were induced by adding 0.3 mM isopropyl-β-D-thiogalactopyranoside (IPTG) for 12 h at 16 °C. Cells were pelleted by centrifugation and resuspended with buffer A [(50 mM Tris (pH 8.0), 200 mM NaCl, and 1 mM PMSF]. The cells were lysed by high-pressure homogenizer at 600-800 bar and then centrifuged at 17,000 rpm for 50 min. Each soluble fraction was passed over a Ni-NTA column (Novagen) and eluted with buffer containing 25 mM Tris (pH 8.0), 200 mM NaCl, and 200 mM Imidazole. Subsequently, the eluates containing recombinant proteins with 10× His-MsyB tag were further purified by an anion-exchange column (Source-15Q; GE Healthcare). The 10×His-MsyB tag was cleaved by TEV protease at 4 °C overnight and removed by an anion-exchange column. Untagged recombinant proteins were then concentrated and further purified by size-exclusion chromatography (Superdex 200 Increase10/300 GL; GE Healthcare) in buffer containing 20 mM Tris (pH 8.0), 200 mM NaCl, and 3 mM DTT. The peak fractions of each protein were pooled together and used for gel filtration assay.

### Gel filtration assay

The recombinant proteins purified as described above were subjected to gel filtration assay (Superdex 200 Increase10/300 GL; GE Healthcare) in buffer containing 20 mM Tris (pH 8.0), 200 mM NaCl, and 3 mM DTT. A mixture of the purified LSD1 and GLK1 (or GLK2) proteins was incubated at 4 °C for 1 h before gel filtration. Samples from relevant fractions were applied to SDS-PAGE and visualized by Coomassie blue staining.

### Measuring 5-ALA synthesis rate

The 5-ALA synthesis rate was quantified as previously described (Goslings et al., 2004). CL-grown 16-day-old plants of WT, *flu, lsd1*, and *lsd1 flu* were vacuum-infiltrated for 5 min with an 80 mM levulinic acid (Sigma) solution containing 10mM KH_2_PO_4_ (pH 7.2) and 0.5% (v/v) Tween 20. After 1 hour incubation at room temperature under CL, samples were immediately frozen in liquid nitrogen and then homogenized in 4% (v/v) TCA. The homogenates were lysed at 95 °C for 15 min, cooled on ice for 2 min, and filtrated with 0.45 μm cellulose acetate membrane filters (Sterlitech). The filtrated lysates were neutralized with an equal volume of 0.5 M NaH_2_PO_4_ (pH 7.5). Afterward, ethylacetoacetate (1/5) was added and then the samples were incubated at 95 °C for 10 min. After cooling on ice for 5 min, the extracts were mixed with the same volume of fresh Ehrlich’s reagent [0.2 g *p*-dimethylaminobenzaldehyde (Sigma-Aldrich), 8.4 mL acetic acid, and 1.6 mL 70% (v/v) perchloric acid (Sigma-Aldrich)] and centrifuged at 14,000 g for 5 min at 4 °C. The OD of each supernatant was measured at 553 nm using the NanoDrop 2000 (Thermo Fisher Scientific). The amount of 5-ALA was calculated using a coefficient of 7.45 × 10^4^ mol^-1^ cm^-1^.

### Determining photochemical efficiency

Measurements of photochemical efficiency of PSII (Fv/Fm) were conducted with a FluorCam system (FC800-C/1010GFP; Photon Systems Instruments) containing a CCD camera and an irradiation system according to the instrument manufacturer’s instructions.

### Trypan blue staining

Cell death was determined by trypan blue (TB) staining as described previously (Lv et al., 2019). The plant tissues were submerged in TB staining solution [25% (v/v) phenol, 25% (v/v) glycerol, 25% (v/v) lactic acid, 0.05% (w/v) trypan blue] diluted with ethanol 1:2 (v/v) and boiled for 2 min. After incubating for 16 h on a vertical shaker at room temperature, the non-specific staining was removed using destaining solution (250 g chloral hydrate dissolved in 100 ml H_2_O, pH 1.2). Plant tissues were then kept in 50% (v/v) glycerol before taking images.

### Gene ontology (GO) enrichment analysis

The individual RNA-seq data using the *lsd1* and *glk1 glk2*, as analyzed in Supplemental Figure 5A, were previously published by Li et al. (Lv et al., 2019) and Ni *et al.* (Ni et al., 2017), respectively. The GO enrichment analysis of the selected genes shown in Supplemental Dataset 5 was performed on gprofiler (https://biit.cs.ut.ee/gprofiler) and represented the significantly enriched GO terms in the data set of biological processes (BP) with a significance of *P*-value < 0.05.

### Pigment analysis

The level of Pchlide was measured in 10-day-old plants of WT, *flu, lsd1*, and *lsd1 flu* as described by Goslings *et al*. (Goslings et al., 2004).

## AUTHOR CONTRIBUTIONS

M.L., K.P.L., T.L., V.D., W.X., and C.K. designed the experiments. M.L., K.P.L., T.L., V.D., J.D., and M.S.L. performed the experiments. M.L., K.P.L., T.L., V.D., W.X., and C.K. analyzed the data. C.K. wrote the manuscript with significant contributions from M.L and K.P.L. All authors discussed the results and reviewed the manuscript.

## ACKNOWLEDGMENTS

We thank the Core Facility of Proteomics in Shanghai Center for Plant Stress Biology (PSC) for carrying out mass spectrometry. No conflict of interest declared.

## SUPPLEMENTAL INFORMATION

### SUPPLEMENTAL FIGURES

**Supplemental Figure 1.**
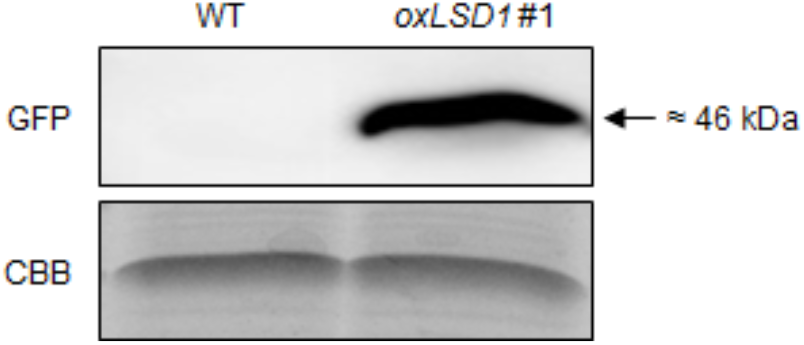
Detection of the LSD1-GFP fusion protein. Total proteins were extracted from two-week-old plants of WT and *35S:LSD1-GFP* transgenic line (*oxLSD1*) grown under CL. The proteins were subjected to an immunoblot assay to detect the LSD1-GFP fusion protein using an anti-GFP antibody. Denaturing gel stained with Coomassie brilliant blue (CBB) was used as a loading control.

**Supplemental Figure 2.**
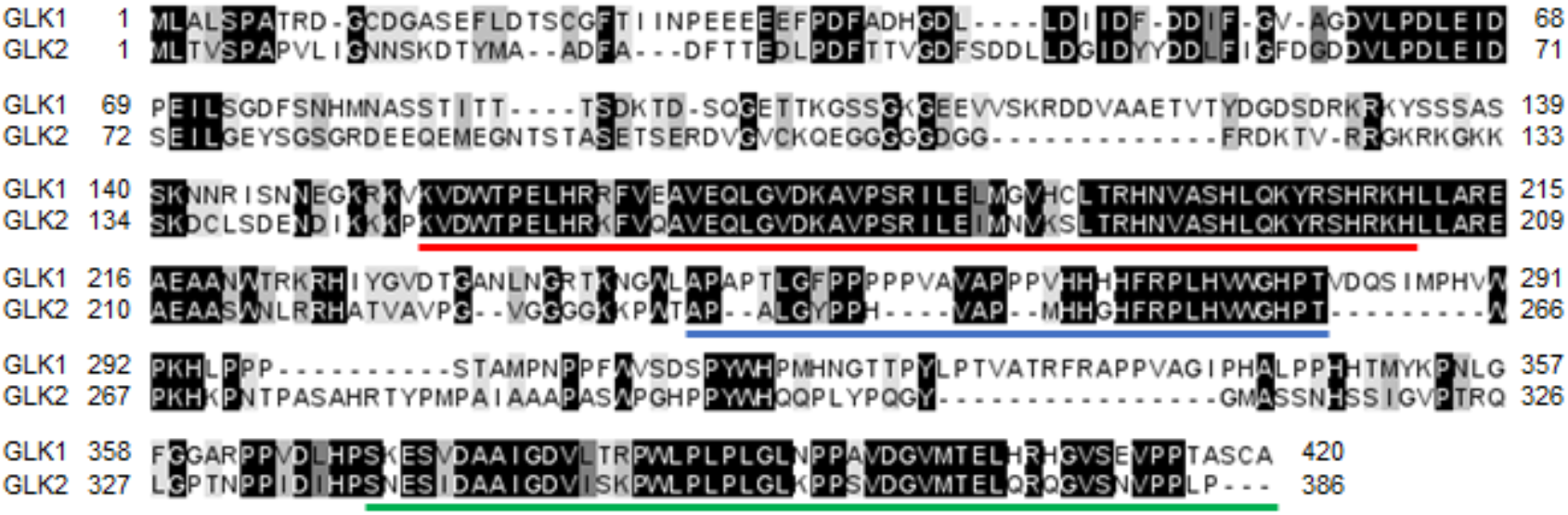
Sequence alignment of Arabidopsis GLK1 and GLK2 proteins. The conserved amino acid sequences were aligned using the Clustal Omega software (https://www.ebi.ac.uk/Tools/msa/clustalo/) before shading with the Jalview program. The boxes with different colors indicate identical and similar amino acids (black: 100%; dark gray: ≥75%; light gray: ≥50%). Red, blue, and green lines indicate region of DNA-binding domain (DBD), proline-rich domain (PRD), and GLK/C-terminal box (GCT-box), respectively.

**Supplemental Figure 3.**
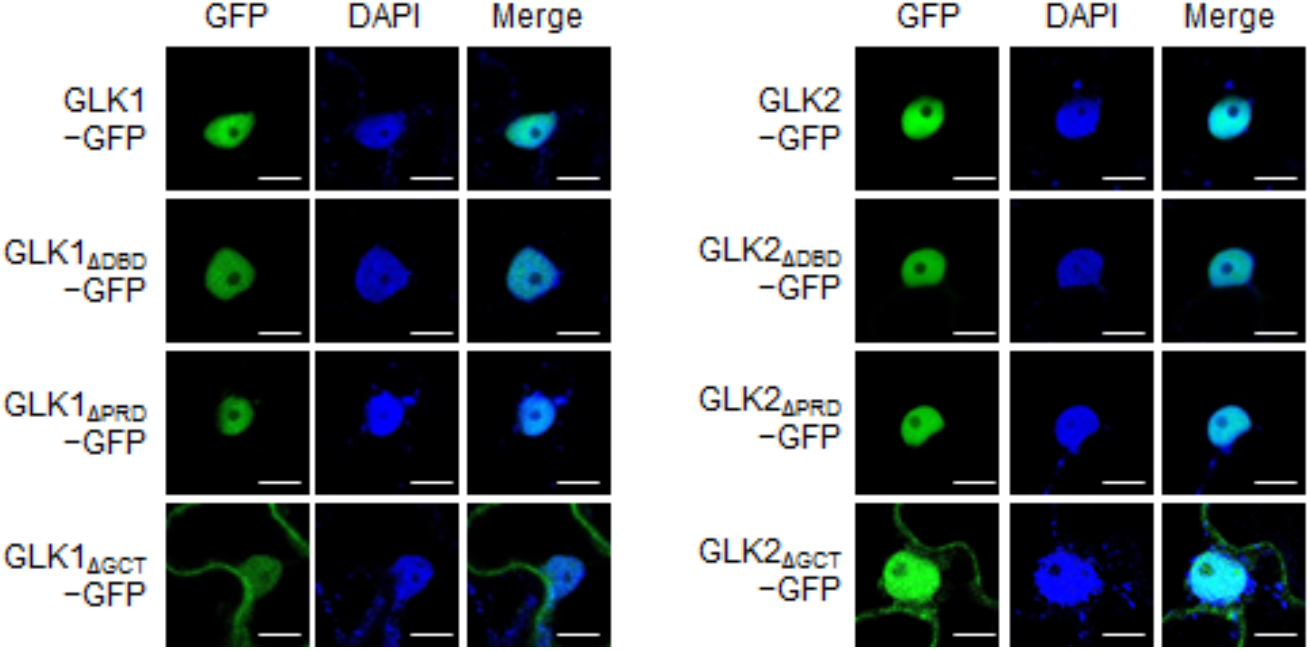
Domain-deleted GLK1 and GLK2 variants localize to the nucleus. Subcellular localization of GFP-tagged intact GLK1/2 and their domain-deleted variants upon transient expression in *N. benthamiana* leaves. DAPI was used to stain the nucleus. All images were taken at the same scale (scale bars: 10 μm).

**Supplemental Figure 4.**
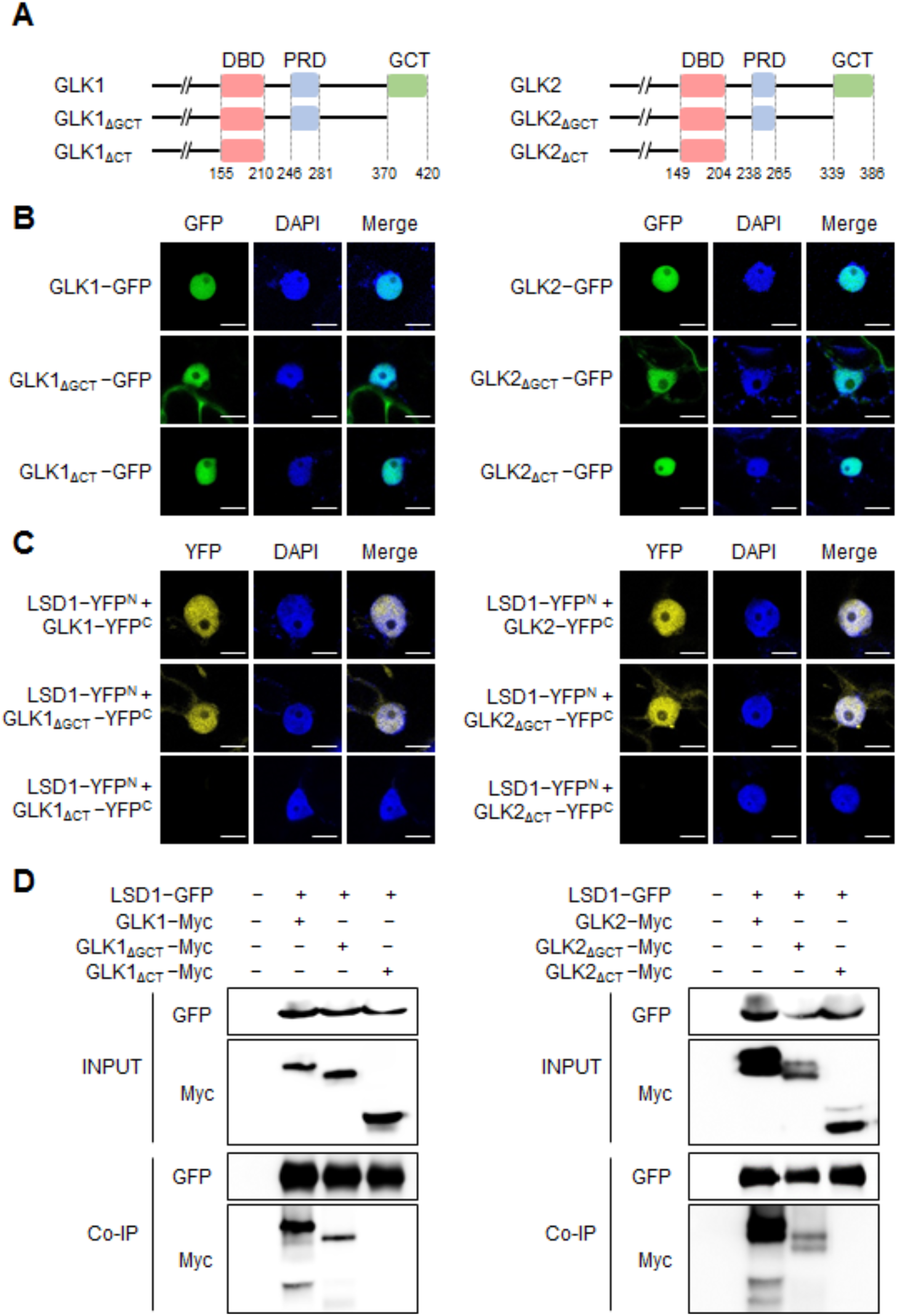
The C-terminal PRD region of GLKs is critical for LSD1-GLKs interaction. **(A)** Schematic diagram of GLK1/2 and their C-terminally truncated variant proteins. **(B)** Subcellular localization of GFP-tagged protein variants **(A)** upon transient expression in *N. benthamiana* leaves. **(C)** BiFC analysis. The intact or truncated variants of GLK1 or GLK2 fused with YFP^C^ were individually coexpressed with LSD1 fused with YFP^N^ in *N. benthamiana* leaves. In **(B)** and **(C)**, DAPI was used to stain the nucleus. All images were taken at the same scale (scale bars: 10 μm). **(D)** Co-IP analyses using *N. benthamiana* leaves coexpressing LSD1-GFP with indicated intact or truncated variants of GLK1 (or GLK2) fused with Myc-tag. GFP-Trap beads were used, and the interaction was evaluated by using Myc antibody.

**Supplemental Figure 5.**
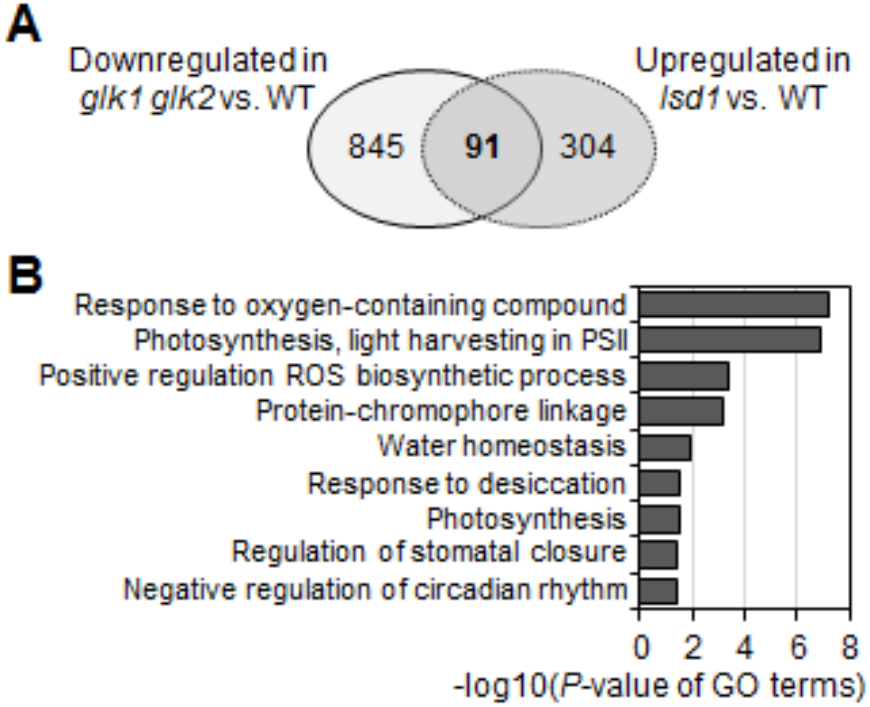
Loss of LSD1 leads to an upregulation of GLK target genes. **(A)** Venn diagram showing the numbers of uncommon and overlapped genes between upregulated genes (395) in 17-d-old *lsd1* (Lv et al., 2019) and downregulated genes (936) in *glk1 glk2* (Ni et al., 2017). **(B)** Gene Ontology (GO) enrichment analysis towards the biological process of the overlapped genes in **(A)**. The GO enrichment analysis was done as described in Methods.

**Supplemental Figure 6.**
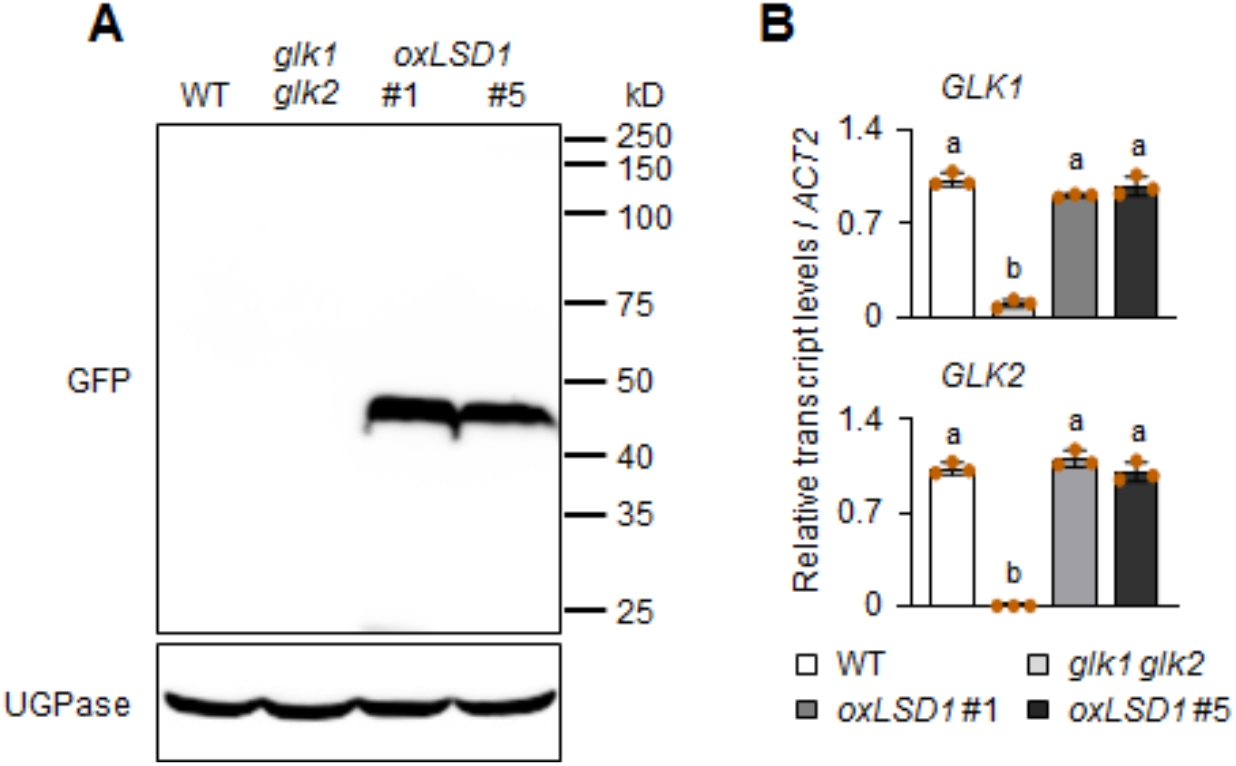
Expression levels of *GLK1* and *GLK2* in *oxLSD1* lines. **(A and B)** Total protein and RNA were extracted from 24-d-old CL-grown plants of WT, *glk1 glk2*, and two independent transgenic lines overexpressing GFP-tagged LSD1 under the control of the CaMV 35S promoter (*oxLSD1* #1 and #5). The proteins were subjected to an immunoblot assay to detect the LSD1-GFP fusion proteins using an anti-GFP antibody **(A)**. UGPase was used as a loading control. The relative expression levels of *GLK1* and *GLK2* were analyzed by RT-qPCR **(B)**. *ACT2* was used as an internal standard. Data are means ± SD (n=3). Lowercase letters indicate statistically significant differences between mean values (P < 0.01, one-way ANOVA with posthoc Tukey’s HSD test).

**Supplemental Figure 7.**
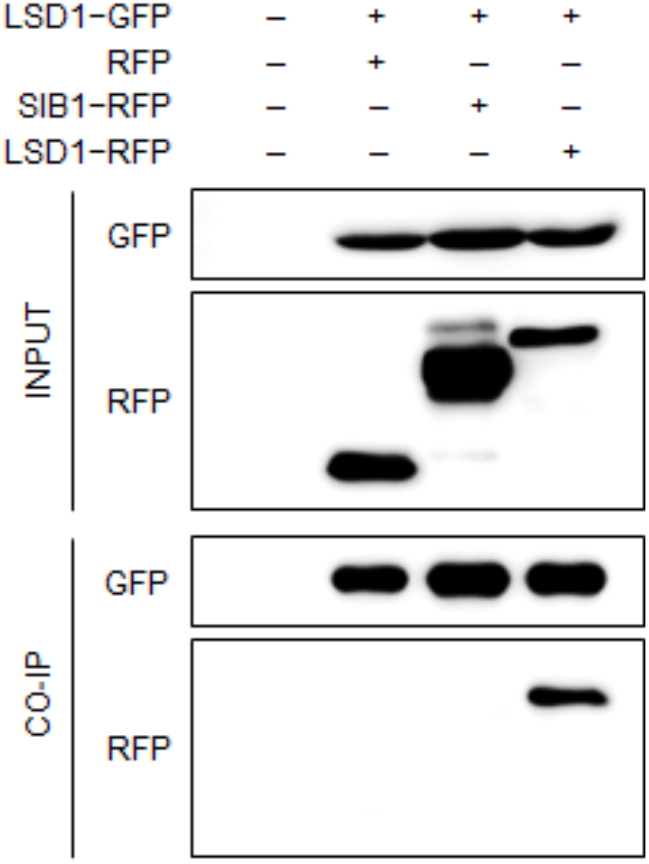
LSD1 does not interact with SIB1 *in vivo.* Co-IP analyses using *Arabidopsis* leaf protoplasts transiently coexpressing LSD1-GFP and SIB1-RFP. LSD1-GFP was also coexpressed with free RFP as a negative control or LSD1-RFP as a positive control. The immunoblot result is one representative of three independent experiments with similar results.

**Supplemental Figure 8.**
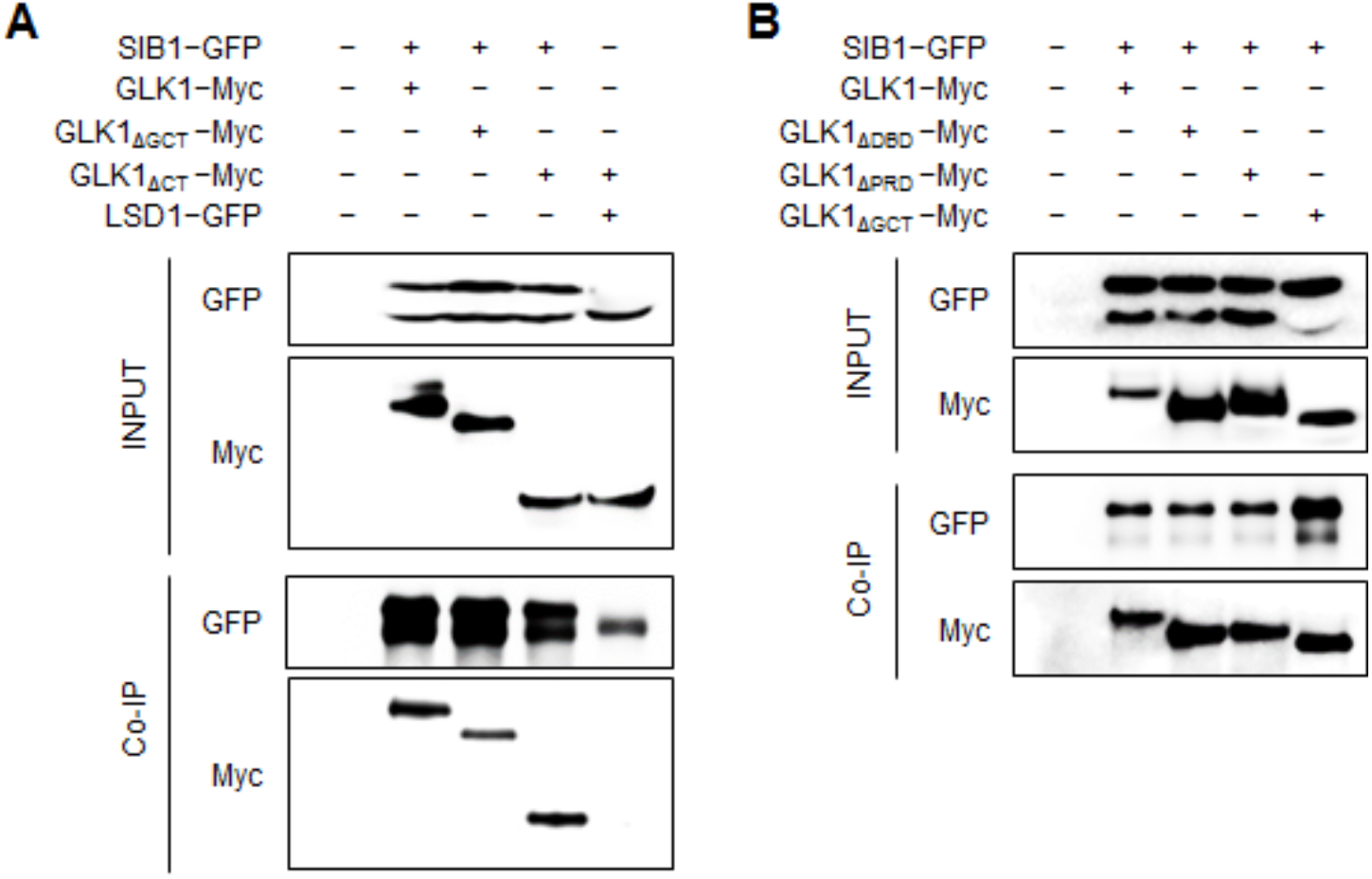
SIB1 interacts with GLK1 through the N-terminal region of GLK1. **(A and B)** Co-IP analyses using *N. benthamiana* leaves transiently coexpressing SIB1-GFP with intact form and C-terminally truncated **(A)**, or domain-deleted variants **(B)** of GLK1 fused with Myc-tag. LSD1-GFP was also transiently coexpressed with GLK1_ΔCT_-Myc as a negative control showing a lack of interaction. Co-IP was performed with GFP-Trap beads, and the interaction was evaluated by using the Myc antibody.

### SUPPLEMENTAL TABLE

**Supplemental Table 1.**
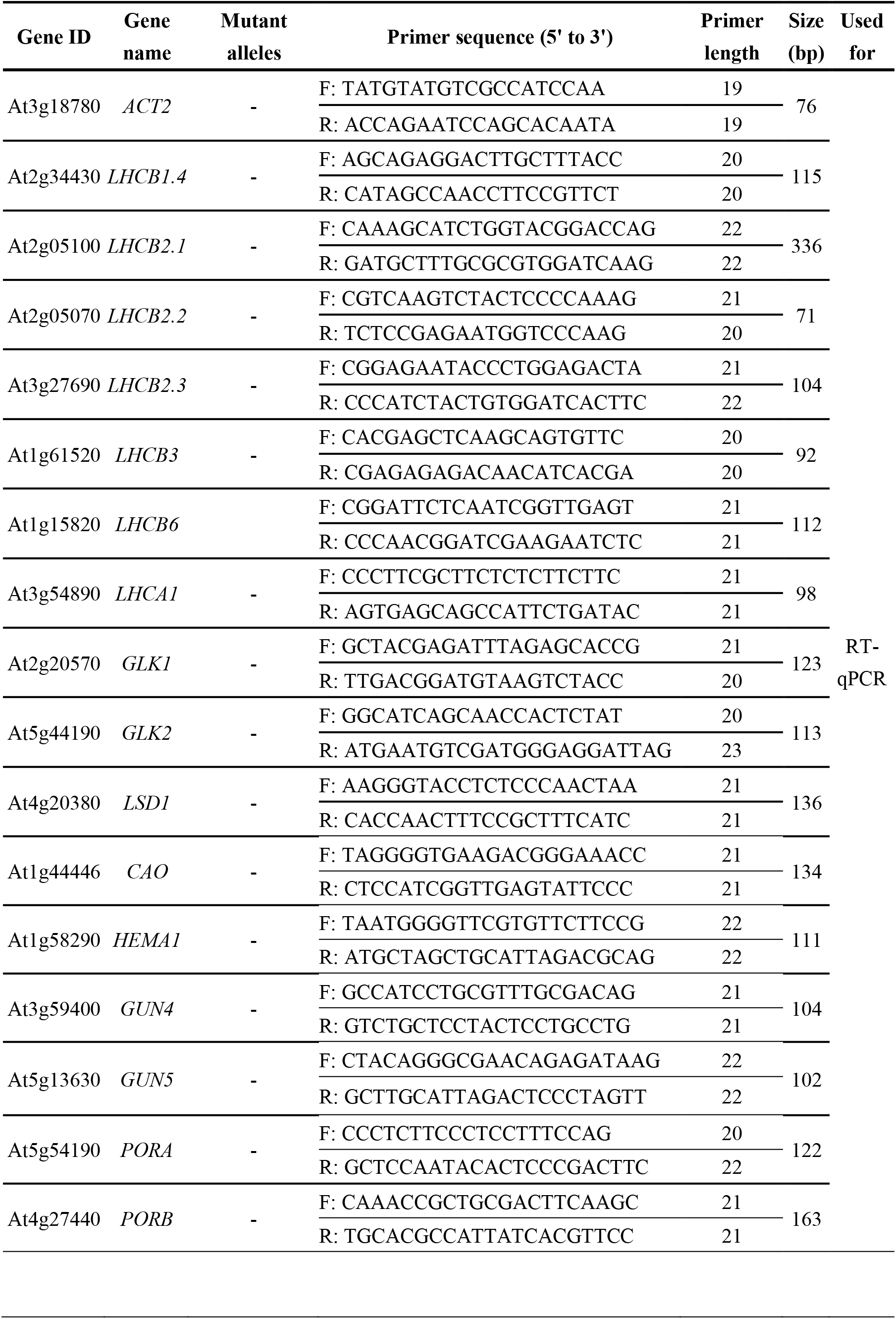
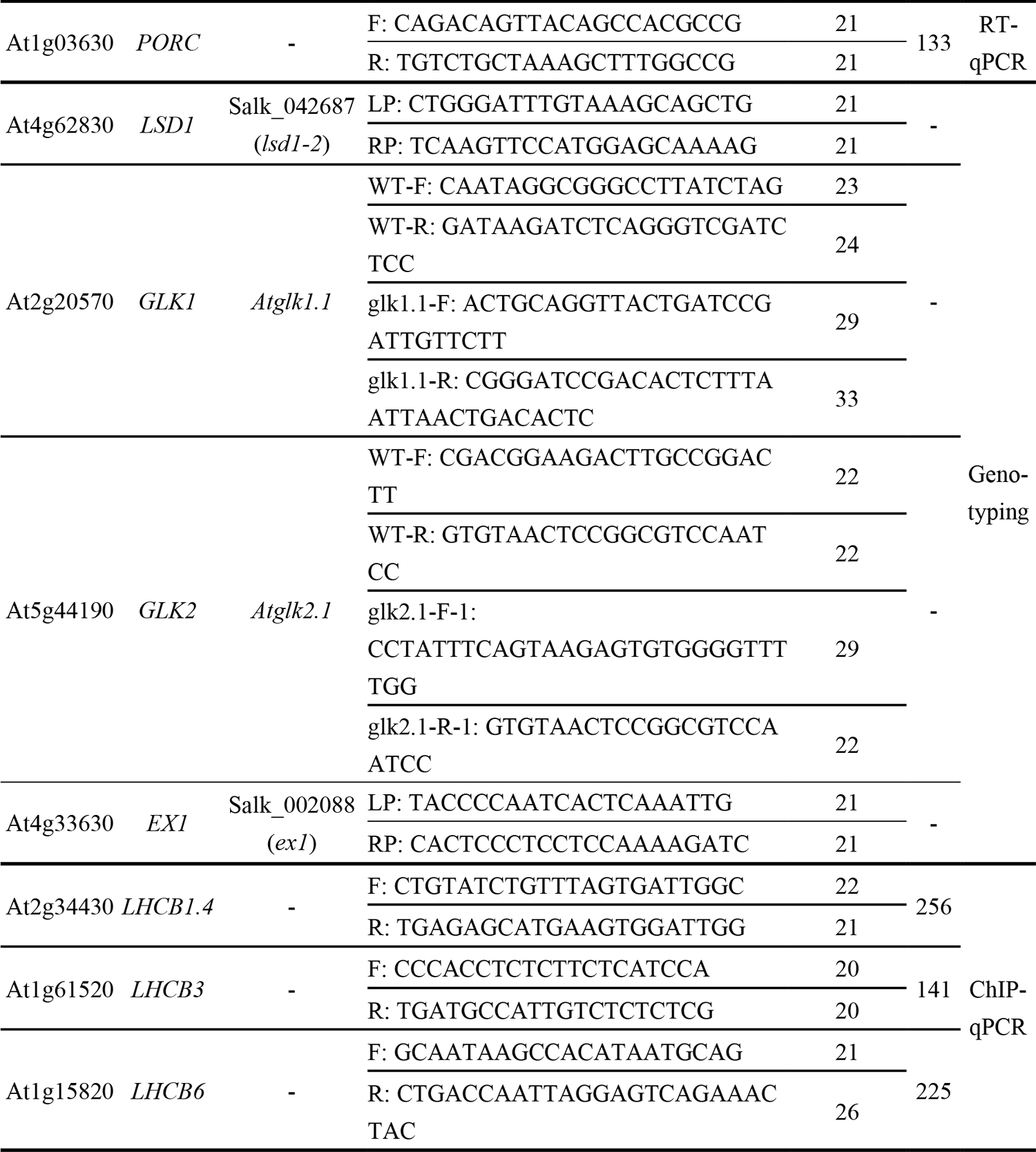
List of primer sets used in this study.

